# Expanded stoichiometric model of chondrocyte metabolism: response to cyclical shear and compressive loading

**DOI:** 10.64898/2026.03.01.708861

**Authors:** Aubrey H. Kimmel, Adrienne D. Arnold, Ayten E. Erdogan, Ronak Kommineni, Erik P. Myers, Breschine Cummins, Ross P. Carlson, Ronald K. June

## Abstract

Cartilage deterioration is a hallmark osteoarthritis, and there is substantial interest in developing strategies for cartilage repair. Cyclical mechanical stimulation has been known for decades to drive synthesis of cartilage matrix proteins. Matrix synthesis requires activation of central metabolism for producing precursors to non-essential amino acids required for protein translation. However, there are gaps in knowledge regarding how mechanical stimuli affect chondrocyte central metabolism. Here, we find that cyclical shear and compression drive differences in chondrocyte central metabolism in a sex-dependent manner. Based on established biochemistry, we developed and tested a stoichiometric model containing 139 metabolites and 172 reactions from central metabolism that includes production of key cartilage matrix proteins. We then used experimental metabolomics data from shear and compressive stimulation of osteoarthritic chondrocytes to constrain this model and ran multiple simulations examining the potential for producing matrix proteins and ATP. Our results show that both shear and compression can stimulate osteoarthritic chondrocyte metabolism in a manner consistent with production of cartilage matrix proteins, with notable differences between male and female chondrocytes. Additionally, and importantly, our simulation results suggest that nitrogen availability is a key limitation to chondrocyte synthesis of matrix proteins. These results are a starting point for using central metabolism of chondrocytes to optimize synthesis of matrix proteins for cartilage repair. For example, increasing glutamine levels in the presence of cyclical compression has potential to increase production of both types II and VI collagen. These strategies have potential for improving cartilage tissue engineering and repair.

## Introduction

Cyclical loading of articular chondrocytes can increase production of cartilage matrix molecules [1]. Expansion of these methods of mechanical stimulation has yielded impressive results with strong potential for cartilage tissue engineering [2]. However, mechanistic details between mechanical stimulation and the degree of matrix production remain unknown. This paper builds on previous mathematical models of chondrocyte metabolism [3, 4] to better understand how cyclical mechanical stimulation affects production of both required metabolic precursors (e.g. ATP) and cartilage matrix molecules (e.g. type VI collagen.)

Stoichiometric modeling is a computational approach often used to quantify intracellular metabolite fluxes by leveraging the structure of biochemical networks [5]. This method is particularly well-suited for studying central energy metabolism, which encompasses glycolysis, the pentose phosphate pathway, and the tricarboxylic acid (TCA) cycle—pathways that generate ATP and precursors for amino acid synthesis (Figure 1) [5]. In the context of articular chondrocytes, stoichiometric models enable the integration of targeted metabolomic data to estimate intracellular fluxes toward ATP and collagen production. Although chondrocytes have traditionally been considered to use primarily glycolytic substrate level phosphorylation for ATP generation, recent evidence suggests they also use respiration and oxidative phosphorylation pathways for ATP generation [6], especially in response to mechanical stimuli. By constructing a stoichiometric matrix that reflects the topology of central metabolism, this study analyzes how physiological loading may alter metabolic activity and support chondrocyte homeostasis. This modeling framework has been successfully applied across diverse biological systems, from microbes to human tissues, and offers a powerful tool for systems-level analysis of chondrocyte function under mechanical stimuli.

**Figure 1:**
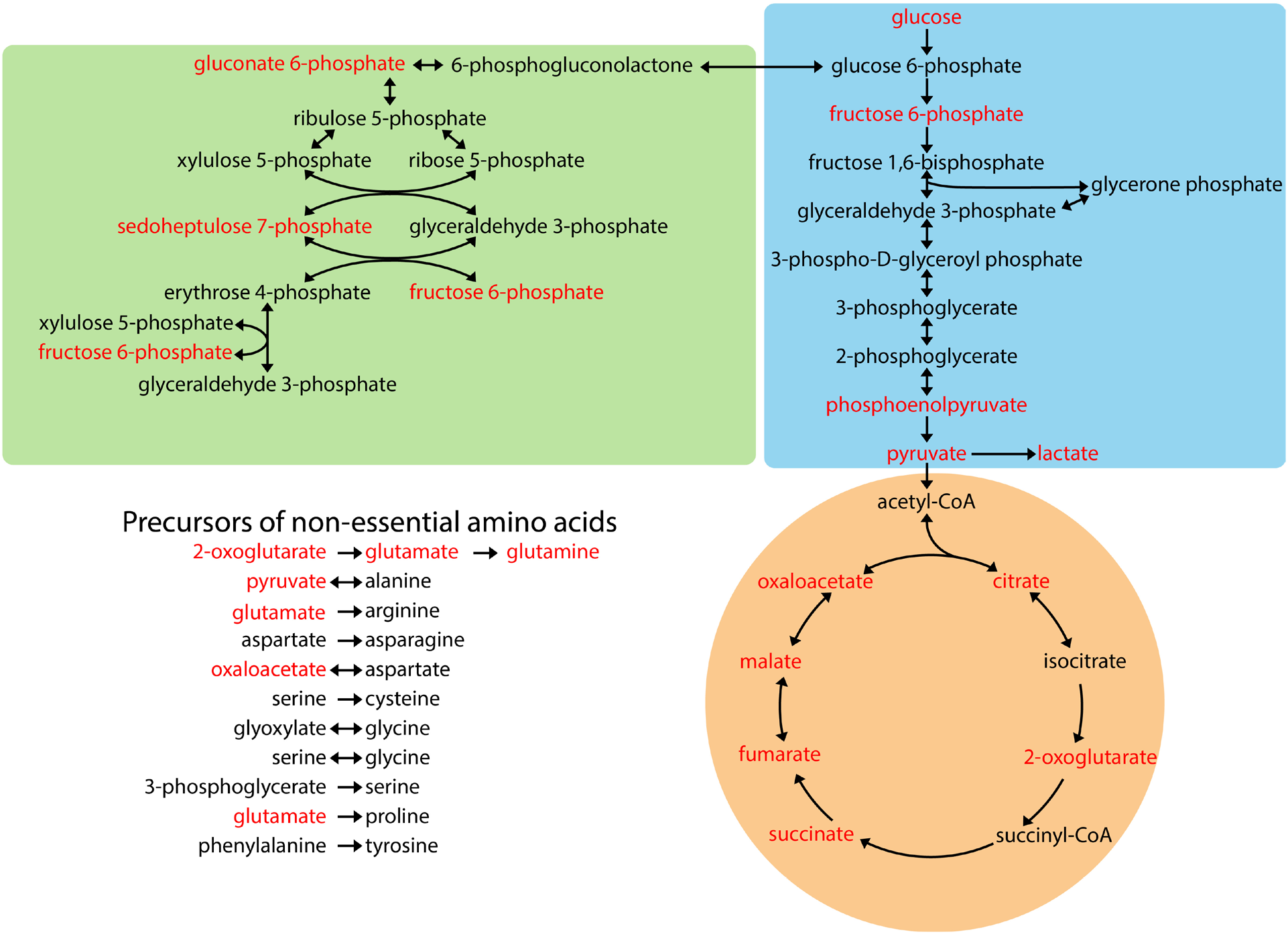
Central carbon metabolism of chondrocytes. Embden-Meyerhof-Parnas glycolysis shown in blue. Pentose phosphate pathway shown in green. Citric acid cycle shown in orange. Abbreviated reactions are included for non-essential amino acid synthesis; these show the carbon backbone precursors of each amino acid. Metabolites in red were measured using liquid chromatography mass spectrometry (Table 1) and are included in the metabolic model as “observed” metabolites which set simulation flux constraints.

**Table 1.** Table of measured (observed) metabolites from 10 human donors, 5 male and 5 female. The data is represented as intervals of the mean plus/minus one standard deviation.

|  | Male |  |  |  | Female |  |  |  |
| --- | --- | --- | --- | --- | --- | --- | --- | --- |
|  | 15C | 15S | 30C | 30S | 15C | 15S | 30C | 30S |
| **Oxaloacetic Acid** | -5.82e-02 ± 1.22e-01 | 7.23e-03 ± 1.55e-01 | -1.03e-01 ± 9.05e-02 | -6.05e-02 ± 9.42e-02 | -1.07e-03 ± 6.55e-02 | 7.23e-03 ± 1.55e-01 | -1.34e-02 ± 3.62e-02 | -3.19e-02 ± 6.03e-02 |
| **Fumaric Acid** | -1.07e-03 ± 7.72e-03 | 4.51e-03 ± 1.96e-02 | -1.31e-03 ± 4.05e-03 | -4.11e-03 ± 1.03e-02 | -4.76e-03 ± 6.02e-03 | 4.51e-03 ± 1.96e-02 | 1.10e-03 ± 6.66e-03 | 3.52e-04 ± 4.38e-03 |
| **PEP** | 1.16e-03 ± 5.33e-03 | 1.80e-02 ± 2.27e-02 | -5.46e-04 ± 1.07e-03 | -1.19e-03 ± 1.85e-03 | 2.97e-03 ± 6.89e-03 | 1.80e-02 ± 2.27e-02 | 8.04e-04 ± 5.67e-04 | 6.28e-03 ± 9.73e-03 |
| **Malic Acid** | -2.10e-03 ± 5.89e-03 | -2.13e-03 ± 3.83e-03 | -1.65e-03 ± 3.39e-03 | -1.22e-03 ± 2.97e-03 | -7.52e-03 ± 4.64e-03 | -2.13e-03 ± 3.83e-03 | -4.15e-03 ± 2.28e-03 | -4.16e-03 ± 3.17e-03 |
| **2-Oxoglutaric Acid** | -2.10e-02 ± 5.05e-03 | 1.65e-02 ± 2.86e-02 | -1.39e-02 ± 9.72e-03 | -2.87e-02 ± 3.94e-02 | -6.17e-03 ± 2.98e-02 | 1.65e-02 ± 2.86e-02 | -3.83e-03 ± 1.29e-02 | -7.94e-03 ± 1.13e-02 |
| **6-P-gluconate** | 9.53e-04 ± 1.85e-03 | 1.05e-03 ± 1.32e-03 | 1.17e-03 ± 3.18e-03 | 9.82e-04 ± 8.76e-04 | 3.49e-04 ± 5.67e-04 | 1.05e-03 ± 1.32e-03 | 4.86e-04 ± 1.37e-03 | -5.01e-05 ± 8.04e-04 |
| **Citric Acid** | -3.19e-04 ± 4.62e-03 | 3.05e-02 ± 4.96e-02 | 3.62e-04 ± 4.20e-03 | 1.57e-03 ± 2.29e-03 | 3.61e-03 ± 9.33e-03 | 3.05e-02 ± 4.96e-02 | 1.81e-04 ± 4.45e-03 | -1.67e-03 ± 2.84e-03 |
| **Sedoheptulose-7-P** | -2.75e-04 ± 1.57e-03 | -1.55e-03 ± 5.80e-03 | -6.50e-04 ± 1.11e-03 | 6.70e-03 ± 1.30e-02 | 2.57e-03 ± 1.74e-02 | -1.55e-03 ± 5.80e-03 | -1.71e-03 ± 1.56e-03 | -2.60e-03 ± 1.34e-03 |
| **Succinic Acid** | 4.48e-03 ± 3.50e-01 | -2.14e-02 ± 1.53e+00 | 1.87e-01 ± 5.27e-01 | 5.45e-02 ± 3.56e-01 | -1.80e-01 ± 1.03e+00 | -2.14e-02 ± 1.53e+00 | 2.24e-02 ± 4.08e-01 | 2.96e-01 ± 2.60e-01 |
| **Fructose-6-P** | 1.41e-02 ± 3.47e-02 | 2.95e-02 ± 5.93e-02 | 8.50e-03 ± 2.49e-02 | -1.43e-02 ± 1.70e-02 | 1.96e-02 ± 4.88e-02 | 2.95e-02 ± 5.93e-02 | 1.79e-03 ± 8.73e-03 | 3.17e-03 ± 1.08e-02 |
| **Glutamic Acid** | -5.88e-02 ± 5.39e-02 | -1.90e-02 ± 9.07e-02 | -3.99e-02 ± 3.11e-02 | -2.22e-02 ± 2.03e-02 | -3.07e-02 ± 9.78e-02 | -1.90e-02 ± 9.07e-02 | -1.98e-02 ± 5.75e-02 | -2.31e-02 ± 4.82e-02 |
| **Lactic Acid** | -7.12e-01 ± 8.94e-01 | -3.46e-01 ± 4.98e-01 | -2.52e-01 ± 3.22e-01 | -4.41e-01 ± 5.46e-01 | -9.10e-01 ± 8.24e-01 | -3.46e-01 ± 4.98e-01 | -1.52e-01 ± 4.03e-01 | -6.10e-01 ± 4.00e-01 |
| **Glutamine** | -5.77e-03 ± 2.21e-02 | -1.08e-02 ± 6.10e-03 | -1.36e-02 ± 8.96e-03 | -2.93e-02 ± 4.39e-02 | -1.78e-02 ± 1.11e-02 | -1.08e-02 ± 6.10e-03 | -7.04e-03 ± 7.59e-03 | -1.15e-02 ± 8.07e-03 |
| **Fructose/ Glucose** | -2.14e-01 ± 1.30e+00 | -2.53e+00 ± 1.82e+00 | -9.96e-01 ± 6.64e-01 | -8.55e-01 ± 1.48e+00 | -4.16e+00 ± 3.55e+00 | -2.53e+00 ± 1.82e+00 | -2.42e+00 ± 1.56e+00 | -3.01e+00 ± 1.19e+00 |
| **Pyruvic Acid** | -2.83e-01 ± 3.35e-01 | -2.86e-01 ± 6.12e-01 | -1.38e-01 ± 1.49e-01 | -1.50e-01 ± 1.72e-01 | -4.71e-01 ± 6.82e-01 | -2.86e-01 ± 6.12e-01 | -2.14e-01 ± 3.18e-01 | -2.80e-01 ± 3.13e-01 |

Prior studies used several *in silico* modeling approaches to understand how chondrocyte metabolism responds to mechanical stimulation. Salinas *et al* showed that cyclical compression synchronized metabolite profiles of central metabolism and compression-induced alterations in the TCA cycle in SW1353 chondrocytes [4]. Further analyses in primary human chondrocytes from joint replacement patients found that cyclical compression can increase production of the metabolite precursors for non-essential amino acids with substantial donor-dependent variation [3]. Additional studies have incorporated glycosaminoglycan biosynthesis when studying intervertebral disc cells [7]. This study builds expands these prior models and integrates them with new experimental data to better understand chondrocyte metabolism in response to applied mechanical stimulation.

The objective of this study was to use stoichiometric modeling and published metabolomics data to analyze how cyclical mechanical loading affects central metabolism and associated production of amino acid precursors and collagen. Toward this end, we first expanded a previous stoichiometric model [3, 4] which now considers central metabolism, respiration, lactic acid fermentation, amino acid uptake and synthesis, synthesis of key cartilage matrix molecules including types II and VI collagen, as well as synthesis of aggrecan. We then tested *in silico* model fidelity by predicting experimentally established relationships between ATP production and O_2_ levels. Next, we integrated previously published experimental metabolomics data from primary chondrocytes [8] with the model to quantify the capacity of chondrocytes to produce ATP, types II and VI collagen, and aggrecan. Finally, we used the computational model and experimental data to identify metabolites that potentially limit the synthesis of collagen and aggrecan and are therefore promising intervention targets. These data provide insight into how cyclical mechanical stimulation drives changes in central metabolism and how these changes are predicted to affect matrix protein production. Application of these data and results may be useful for maximizing neomatrix production for tissue engineering and cartilage repair through rational intervention strategies such as targeted amino acid supplementation.

## Materials and Methods

### Experimental Data

This study used experimental data from a previous study [8]. Primary human chondrocytes from joint replacement tissue from stage IV osteoarthritis patients (n=5 male and n=5 female) were extracted and cultured for 1 passage using established methods [9]. Chondrocytes were then encapsulated in 4.5% wt/vol high-stiffness agarose [10] at 500,000 cells per gel and equilibrated in cultured for 24 hours. 1 hour before mechanical stimulation, media was replaced with phosphate buffered saline.

Samples from each donor were randomized into five experimental groups: 15 minutes of cyclical shear strain (15S), 15 minutes of cyclical compressive strain (15C), 30 minutes of cyclical shear strain (30S), 30 minutes of cyclical compressive strain (30C), or unloaded controls. Mechanical strain was applied using a sinusoidal waveform at 1.1Hz with a mean strain of 5% and an amplitude of 2.5% after a 15-minute equilibration of a 5% compressive prestrain. Immediately after mechanical stimulation, samples were removed and bisected. Metabolites were extracted from one half of the sample, and central metabolites were quantified using targeted LC-MS metabolomic profiling. The accumulation or depletion of metabolite levels were determined by subtracting the value of the unloaded control from the value obtained in each experimental group for each donor. These metabolite data provide insight into whether a given metabolite was produced or consumed during the specific mechanical stimulation (e.g. 30 minutes of cyclical shear strain).

### Stoichiometric Modeling

A core-scale stoichiometric model of a human chondrocyte metabolism was constructed using human metabolic network data from HumanCyc and literature (Figure 1) [11]. The model consists of 139 metabolites and 172 reactions representing glycolysis, the citric acid cycle, the aerobic electron transport chain, lactate fermentation, the pentose phosphate pathway, transport and synthesis of amino acids, and synthesis of type II collagen, type VI collagen, and the aggrecan core protein (Figure 1). Collagen composition for both Col2A1 (type II collagen) and Col6A1 (type VI collagen) was determined using the amino acid sequences from HumanCyc [12]. Aggrecan core composition was determined using the canonical amino acid sequence from UniProt [13] (primary accession P16112). The collagen and aggrecan protein sequences represented in the model reactions were scaled to 100 amino acids for consistency and convenience. Reaction directionality (forward, reverse, reversible) was applied based the thermodynamic information from HumanCyc. The model was constructed in CellNetAnalyzer 2019 and 2023.1. All model reactions satisfied conservation of mass, atoms, and electrons [14]. Flux balance analysis was performed using CobraToolbox with MATLAB R2019b. Plotting was performed using MATLAB R2024b. Metabolic model details can be found in SI 1. The code used to analyze the model can be found in SI 3 and SI 4.

Metabolite exchange between mass balanced and unbalanced compartments was controlled using three classes of reactions (Figure 2): void, observed, and transport. Void reactions denote transport between the balanced extracellular environment and an unbalanced void. Observed exchange reactions incorporate experimental metabolomics data and account for increases or decreases in metabolite pools; these reactions enforce consumption or production of metabolites over the experimental period of cyclical mechanical stimulation. For the void and observed reactions, a negative reaction flux indicates that a metabolite is consumed, while a positive reaction flux indicates that a metabolite is excreted. Transport reactions denote exchange between the balanced extracellular compartment and the balanced intracellular environment. The directions of the transport reactions were based on expected physiological function (e.g., glucose flux into the cell would be positive, and CO2 export would also be positive).

**Figure 2:**
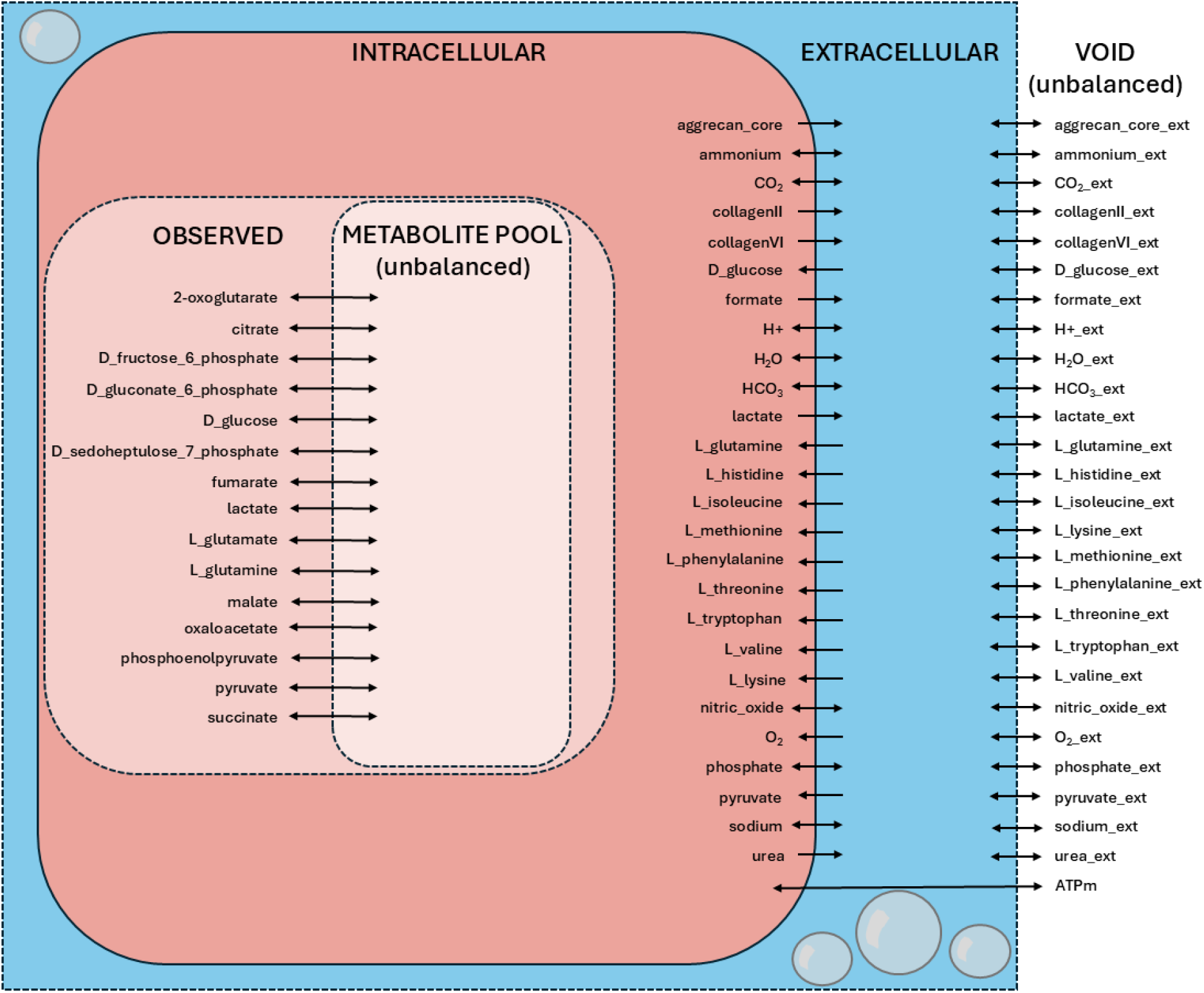
Transport reactions within the metabolic model. The extracellular and intracellular compartments are balanced. The void and observed compartments are unbalanced. Observed reactions incorporate mass spectrometry data. For transport between the void compartment and the extracellular compartment and between the metabolite pool compartment and the observed compartment, negative fluxes = consumption (entering a balanced compartment) and positive fluxes = production (leaving a balanced compartment). For transport between the extracellular and intracellular compartments, reaction directionality is based on expected physiological function. Arrow direction indicates reversibility (bidirectional arrows) or irreversibility (unidirectional arrows). Additional optimization constraints can restrict reversible reactions to irreversible reactions.

### Description of Simulations

Simulations were performed on each of eight groups, where a group was defined based on a combination of an experimental condition (15C, 15S, 30C, or 30S) and sex (M or F). Each simulation consisted of a flux balance optimization [5] given a set of metabolite flux constraints and a maximization objective (e.g. ATP, collagen II, collagen VI, or aggrecan core). Two types of simulations were performed based on applied flux constraints. First, theoretical network properties were quantified considering the composition of the growth medium, namely glucose. These simulations tested model functioning including ATP yield on substrates. The second type of simulation incorporated experimental mass spectrometry data as flux constraints through the ‘observed’ reactions. These simulations were constrained by experimental data and provide insight to the actual distributions of metabolic fluxes of the chondrocytes.

Multiple experimental observations permitted the estimation of a 95% confidence interval for each intracellular metabolite flux and each group. During simulations, the observed fluxes were independently sampled uniformly within these confidence intervals and then applied as flux constraints during flux balance analysis (FBA). Observed fluxes were reasonable estimates of the resources available to the cell. However, there were unmeasured intracellular metabolites due to technical limitations [5]. If a sampled set of observed fluxes resulted in the net secretion of carbon, electron, or nitrogen (i.e. violation of conservation relationships), the set was discarded, and new values were sampled until net consumption was achieved. This resampling (with results shown in Figure 3 reflecting disparate variances between male and female data) minimized infeasible optimization problems due to insufficient carbon, electron, or nitrogen sources. Non-observed metabolites were left unconstrained up to 10,000 fmol/min/cell to avoid biasing predictions and were assigned biologically motivated irreversibility where indicated in Figure 2. O_2_ uptake was constrained to an upper bound of 10 fmol/cell/min based on experimental data [8]. Distributions of reaction rates were compared between groups using the Kolmogorov-Smirnov test with Bonferroni corrections for multiple comparisons.

**Figure 3:**
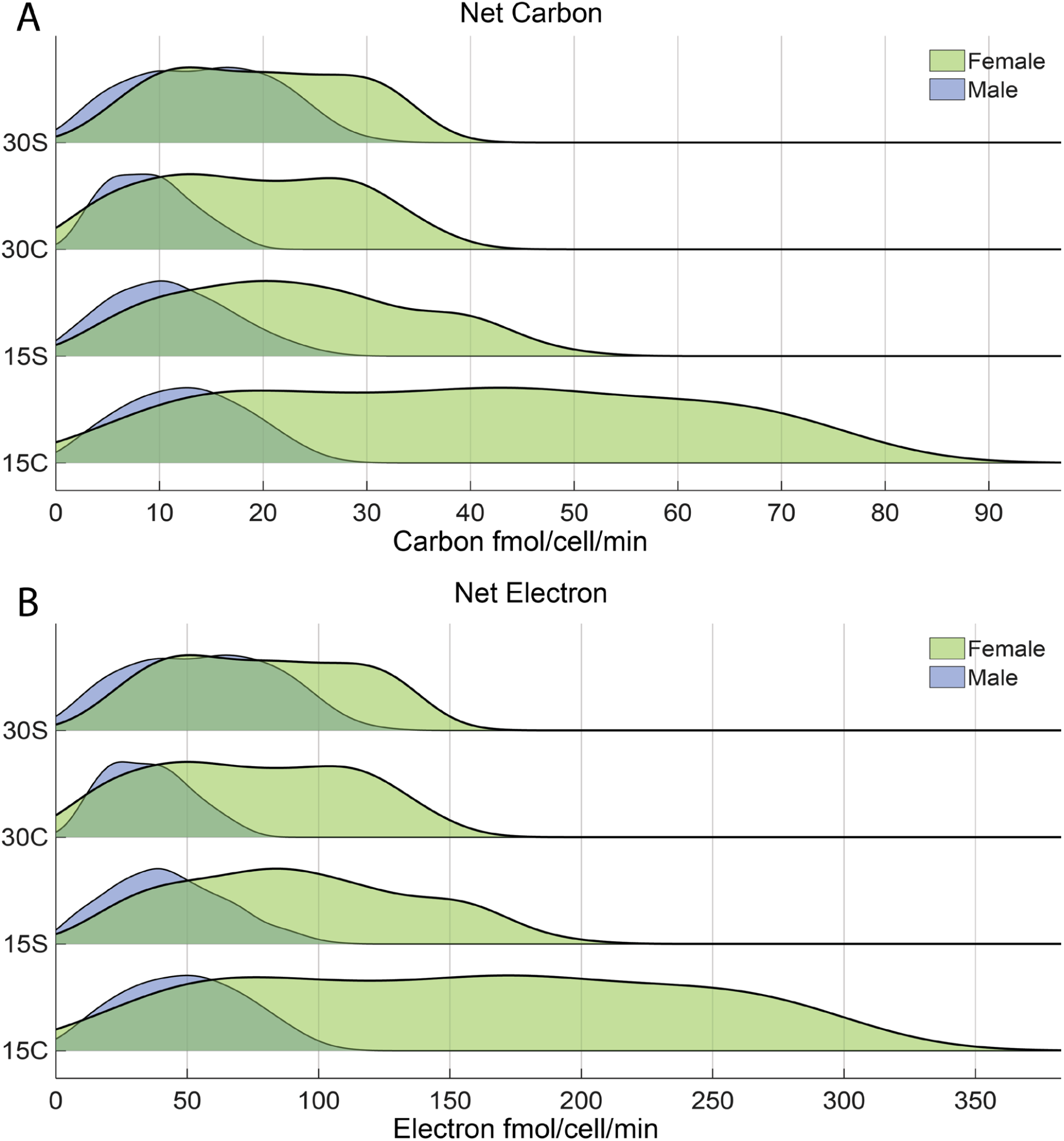
Distributions of net carbon and electron input (fmol/cell/min) for each permutation of time, condition, and sex after checking for net consumption (instead of production). Male data in blue and female data in green. Male and female data are further split into four experimental groups from bottom to top: 15 minutes of compression (15C), 15 minutes of shear (15S), 30 minutes of compression (30C), and 30 minutes of shear (30S). These net carbon and electron inputs were calculated from samples of the observed metabolite source distributions. Female donors appear to have a wider distribution of carbon and electron input than males.

## Results

Model fidelity including carbon and electron balancing and cellular energy representation was verified by (1) checking ATP and lactate yields on glucose and by (2) checking ATP yield as a function of available O_2_ in the presence of glucose. As expected, ATP yields per glucose were 32 mol/mol in aerobic conditions and 2 mol/mol in anaerobic conditions. Also as expected, maximum lactate yields on glucose were 2 mol/mol (results in Supplemental Table S1). ATP yield decreased with decreasing O_2_, as ATP generation shifted from oxidative to substrate-level phosphorylation (results in Supplemental Table S2). We concluded that the model accurately captures essential aspects of central carbon metabolism.

Experimentally measured metabolite fluxes were applied to the model as flux constraints, and the potential to produce ATP, type II or VI collagen, or aggrecan core protein was evaluated. 1000 simulations for each group and the four different maximization objectives were utilized. Two types of predictions were run. First, we ran predictions where observed metabolite fluxes were the only permitted substrates (Figures 4 and 5). Next, we ran predictions where, in addition to the observed metabolites, additional glutamine and ammonium were provided to the system (Figure 6). For each type of prediction, production of ATP, collagen II and VI, and aggrecan core protein were maximized. In Figure 4, the observed metabolite fluxes were the only permitted substrate fluxes. Simulation results for the male metabolomics data are in blue and for the female samples are in green. Each experimental condition is represented by a separate distribution from left to right as 15 minutes of compression (15C), 15 minutes of shear (15S), 30 minutes of compression (30C), and 30 minutes of shear (30S). Each point in each distribution represents the maximum product flux (fmol/cell/min) from each simulation. The median maximum product flux is denoted by the black bar.

**Figure 4:**
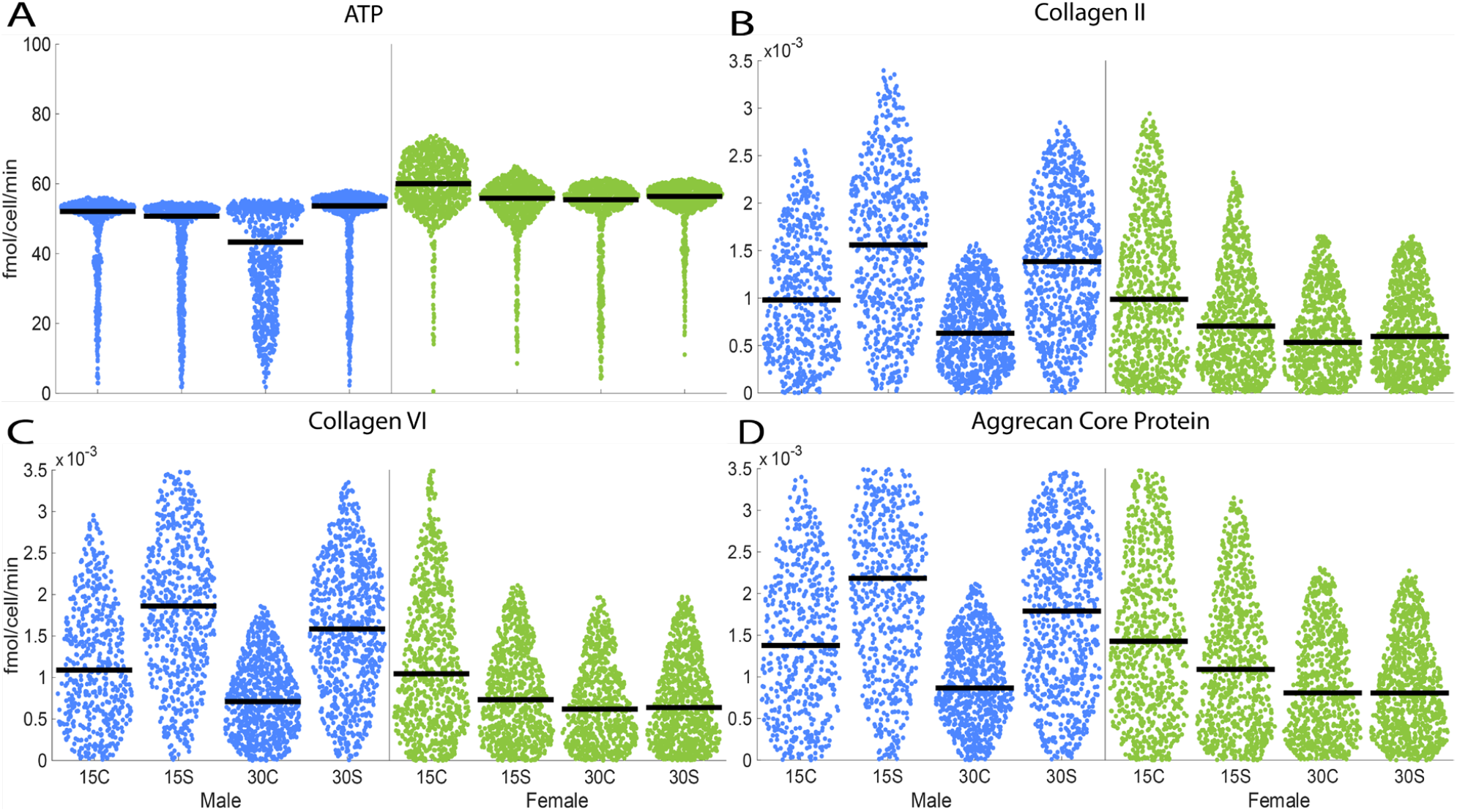
Distributions of maximizing fluxes (fmol/cell/min) over 1000 simulations per condition and optimization criteria. Each point in each distribution for each group represents the maximum flux through the production reaction for the maximization objectives (A) ATP, (B) collagen II, (C) collagen VI, or (D) aggrecan core protein. Only observed metabolic measurements were allowed as sources. The median maximum flux is denoted by the black bar in each plot. In each panel, each experimental condition is represented by a separate distribution from left to right as 15 minutes of compression (15C), 15 minutes of shear (15S), 30 minutes of compression (30C), and 30 minutes of shear (30S), with male data in blue and female data in green.

**Figure 5:**
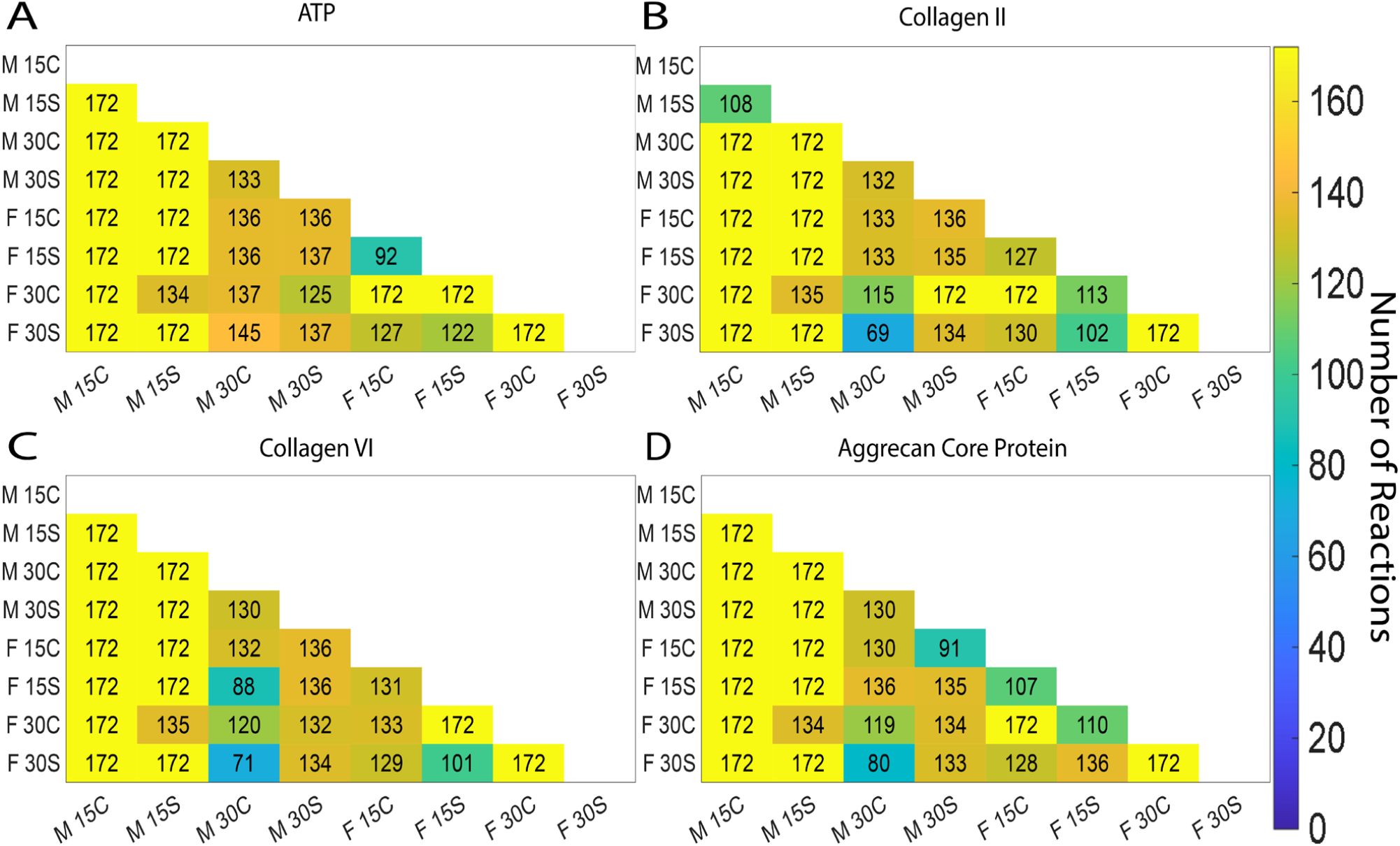
Pairwise tallies of significantly different numbers of reactions between groups with maximization objectives (A) ATP, (B) collagen II production, (C) collagen VI production, and (D) aggrecan core protein. Each cell contains the number of significantly different maximizing flux reactions (from 1000 simulations) out of 172 total reactions between the row and column groups. Statistical significance was determined via the Kolmogorov-Smirnoff test with a p-value adjusted by the Bonferroni Correction Method.

**Figure 6:**
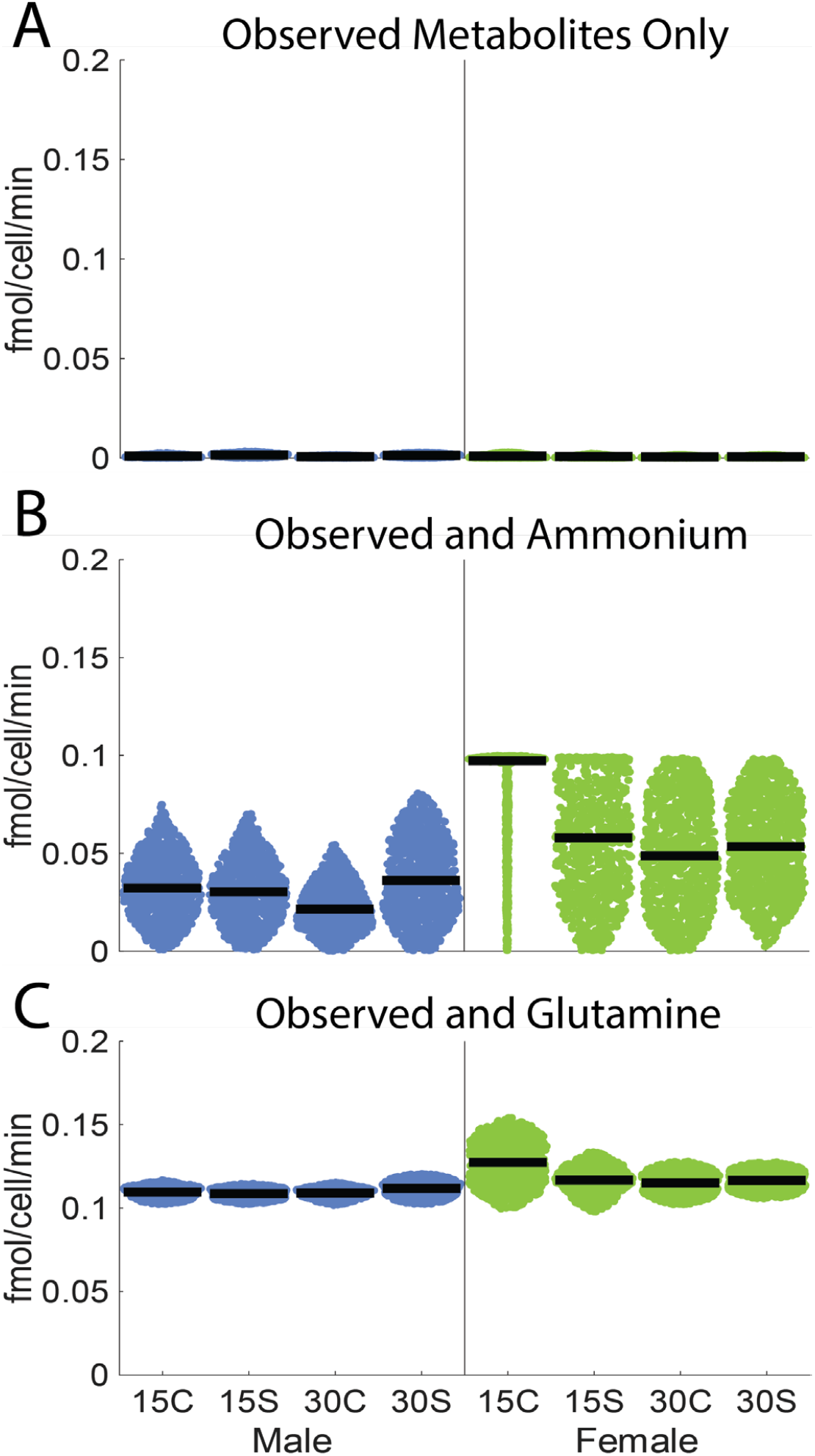
Distributions of type II collagen maximizing fluxes under different uptake allowances. (A) observed metabolite sources only, (B) ammonium uptake allowed, and (C) glutamine uptake allowed. Glucose and pyruvate uptake allowed were also tested, but the results were similar to the observed metabolite sources only plots (see Supplemental Figure S2). ATP, collagen VI, and aggrecan core protein were also simulated and showed qualitatively similar results (see Supplemental Figures S1, S3, S4). These data suggest that nitrogen availability may limit production of type II collagen.

The predicted maximum ATP fluxes were similar across sex and condition in simulations where observed metabolomics data were the only permitted substrates (Figure 4A). While the median fluxes were similar across sex and condition, the differences between groups’ overall distribution shapes and spread reflected the differences in experimental metabolite data (Table 1). In the 30C male group, the flux distributions were more variable, with a wider tail toward low ATP fluxes. For the female samples, most of the distributions looked similar across conditions, with 15C having a slightly higher median and greater spread than the others. The tails did not extend as close to zero as the distributions for the male samples. Males had the lowest median ATP production in the 30C groups, whereas females had the highest median ATP production in the 15C group.

Collagen and aggrecan core protein fluxes were maximized in Figures 4B-D. These data had higher relative variance between groups while there were fewer qualitative differences between simulation objectives (collagen II, collagen VI, aggrecan). The most noticeable difference for male distributions between groups was that mechanical stimulation by cyclical shear showed higher median collagen and aggrecan core protein yield than the cyclical compression for both 15 and 30 minutes. The 15S group was slightly higher than the 30S group, whereas the 30C group was higher than the 15C group. The distributions for 15C and 15S for the female samples had slightly higher median collagen and aggrecan core protein yield than 30C and 30S. The highest median production occurred in the male distributions under shear conditions. We quantified these observations by testing for statistical significance in the difference between flux distributions per group per objective using the Kolmogorov-Smirnoff test with a p-value adjusted by the Bonferroni Correction Method and summing up the number of statistically different individual reaction distributions. The higher the sum, the greater number of statistically different reaction distributions there were in that specific comparison. Most flux distributions within an objective class (ATP, collagen II, collagen VI, or aggrecan core protein) were significantly different, with a few exceptions (see Supplemental Figure S1).

Each simulation resulted in fluxes for all 172 model reactions. We analyzed the statistics of the fluxes across all 172 reactions in the metabolic model. The male and female chondrocytes responded to cyclical shear and compression differently. Figure 5 summarizes comparisons with each cell containing the tally of statistically different reaction distributions (out of 172) when comparing the row group to the column group. Group differences were roughly the same across type II collagen, type VI collagen, and aggrecan core protein simulations. The groups that were the most similar (i.e., had the fewest significantly different flux distributions) were male 30C and female 30S, whereas the male 15C and 15S groups were the most distinguishable from other groups.

The systems-based analysis was used to predict which anabolic resource (carbon, nitrogen, or ATP) was limiting synthesis of product. 1000 simulations were performed where the observed metabolite fluxes were augmented with either glutamine or ammonium. These simulations were performed for each of the four maximization objectives across all groups with the upper bound of O_2_ flux capped at 10 fmol/cell/min. Data for type II collagen is shown in Figure 6. A baseline scenario was set where only the observed metabolite fluxes were permitted with no additional nitrogen or carbon sources present. Results for the other objectives, ATP, collagen VI, and aggrecan core protein, can be found in the supplemental data (Supplemental Figures S2-S5). It should be noted that both collagen VI and aggrecan core protein had similar results to collagen II, while the ATP simulations had no noticeable difference from baseline (Figure 6A).

Significantly higher type II collagen fluxes were possible when an additional nitrogen source was provided to the model (Figures 6B-C). Therefore, we propose nitrogen source is limiting protein synthesis under the tested conditions. When carbon sources glucose or pyruvate were added to the simulations, there was no discernible difference in collagen or aggrecan synthesis from Figure 6A (see Supplemental Figures S2-S5). Carbon source supplementation did enhance the potential for ATP production which was expected. In short, these simulations suggest that production of collagen and aggrecan are not carbon limited.

The addition of glutamine to the simulations resulted in higher protein fluxes than just ammonium (Figure 6B-C). Simulations of both male and female samples showed approximately two orders of magnitude more protein production in the presence of glutamine compared to base simulations, with females increasing slightly more than the males. While also resulting in substantially higher protein production, the addition of ammonium to simulations did not result in the same increase as simulations with additional glutamine. Glutamine can serve as both a nitrogen and carbon source, whereas ammonium is strictly a nitrogen source like resulting in carbon limitation. Adding additional glutamine to the simulations also resulted in less flux variability than in the augmented ammonium simulations. While the male flux distributions are similar across groups, the female flux distributions show higher synthesis rates for type II collagen with 15 minutes of compression for both nitrogen sources. The female flux distributions also have higher variance, while the males are distributed more tightly about the median. This is likely due to the larger variances in the measured metabolomics data from the female samples (Figure 3). For the female samples, 15 minutes of compression resulted in the highest collagen II production flux with the other three conditions showing similar distributions, whereas the results for males were roughly the same across all four conditions and times (Figure 6).

## Discussion

The cyclical loading of chondrocytes has long been shown to increase matrix synthesis [1]. Previous studies found that cyclical compression induces proteoglycan and collagen synthesis [15] as well as the depletion of ATP, which is needed for protein synthesis [16]. Cyclical fluid shear of monolayer chondrocytes results in synthesis and release of both inflammatory mediators as well as proteoglycans [17]. Stoichiometric modeling provides systems-level information regarding how mechanical stimuli can affect the potential of the central metabolism to generate cellular energy and synthesize protein. For example, in SW1353 chondrocytic cells, compression was predicted to induce time-dependent changes in central metabolism consistent with protein synthesis [18]. Furthermore, in primary human chondrocytes, cyclical compression induced metabolic changes consistent with production of metabolic precursors for synthesis of non-essential amino acids [3].

In this study, we used computational modeling of metabolism to analyze how mechanical stimulation by cyclical shear and compression affected chondrocyte central metabolism including production of key amino acid precursors, ATP, and simulated production of key matrix proteins. Using stoichiometric modeling and flux balance analysis, we examined how differences in male and female metabolomic data resulted in potential differences in production of ATP, collagen II, collagen VI, and aggrecan core protein, important components in the production of collagen matrix molecules.

Simulations using only observed metabolite data as constraints highlighted differences in potential collagen and aggrecan production between male and female cells. While results based on female metabolite data showed small differences in median collagen and aggrecan core protein production between most mechanical simulation groups, the results for males showed noticeably higher median collagen and aggrecan production in shear compared to compression (Figure 4). This observation warrants further investigation into whether chondrocyte mechanotransduction of compression and shear is sex-specific. When the distributions of all 172 reactions in the metabolic model were considered, we predicted many significantly different metabolic fluxes between male and female samples for both shear and compressive mechanical stimulation. The computational representation of metabolism was used to investigate what cellular resources were limiting product synthesis (carbon, nitrogen, ATP). These simulations predicted that both males and females would be able to support higher collagen II fluxes if supplemented with glutamine or ammonium. This suggests the chondrocyte metabolism might be nitrogen-limited, and not carbon- or ATP-limited. The highest increase in collagen II production was observed in the female 15 minutes of compression group, suggesting that physical treatment has the potential to be more effective in females. Studies seeking to maximize matrix production for cartilage tissue engineering should ensure sufficient nitrogen availability. Future studies will evaluate the efficiency of nitrogen sources on matrix synthesis.

There are important limitations to this study. First, only a fraction of the intracellular metabolites were measured, which could affect the simulation results as the accumulation of unmeasured metabolites was assumed to be zero. Expanding the number of measured metabolites will provide additional insight into group similarities and differences. Second, the experimental methods could not distinguish between intracellular and extracellular metabolites. Distinguishing between intracellular and extracellular metabolites has the potential to increase the impact of these results.

## Conclusions

To our knowledge, this is the first study integrating metabolomic data from shear and compressive stimulation of primary osteoarthritic chondrocytes with stoichiometric modeling and flux balance analysis. These data provide preliminary evidence that there are significant differences in the mechanically induced central metabolism between sexes and loading type. Furthermore, simulations suggest that nitrogen limitation may play a role in collagen production in osteoarthritic chondrocytes. Future studies may build on this work to advance neomatrix production for cartilage tissue engineering through metabolite interventions.

## Acknowledgements

This study was supported by research grants from the NIH (NIAMS R01AR073964 and R01AR081489) and NSF (CMMI 1554708). AK was partially supported by NSF DMS-1748883. AK and BC were partially supported by NSF AI Institute in Dynamic Systems under award NSF #2112085.

## CRediT Attributions

Conceptualization: BC, RPC, and RKJ

Formal analysis: AHK, ADA, BC, RPC, and RKJ

Funding acquisition: RKJ and BC

Investigation: AHK, ADA, AEE, EM, and RK.

Methodology: ADA, BC, and RPC.

Supervision: BC, RPC, and RKJ

Validation: AHK, ADA, BC, RPC, and RKJ

Writing – original draft: AHK, ADA, AEE, BC, and RKJ

Writing – review & editing: all authors

## References

[1] Sah, R. L., Kim, Y. J., Doong, J. Y., Grodzinsky, A. J., Plaas, A. H., and Sandy, J. D., 1989, “Biosynthetic response of cartilage explants to dynamic compression,” J Orthop Res, 7(5), pp. 619–636.

[2] Mauck, R. L., Byers, B. A., Yuan, X., and Tuan, R. S., 2007, “Regulation of cartilaginous ECM gene transcription by chondrocytes and MSCs in 3D culture in response to dynamic loading,” Biomech Model Mechanobiol, 6(1-2), pp. 113–125.

[3] Salinas, D., Mumey, B. M., and June, R. K., 2019, “Physiological dynamic compression regulates central energy metabolism in primary human chondrocytes,” Biomechanics and Modeling in Mechanobiology, 18(1), pp. 69–77.

[4] Salinas, D., Minor, C. A., Carlson, R. P., McCutchen, C. N., Mumey, B. M., and June, R. K., 2017, “Combining Targeted Metabolomic Data with a Model of Glucose Metabolism: Toward Progress in Chondrocyte Mechanotransduction,” PLoS One, 12(1), p. e0168326.

[5] Doran, P. M., Morrissey, K., and Carlson, R. P., 2024, Bioprocess Engineering Principles, Academic Press.

[6] Wu, X., Liyanage, C., Plan, M., Stark, T., McCubbin, T., Barrero, R. A., Batra, J., Crawford, R., Xiao, Y., and Prasadam, I., 2023, “Dysregulated energy metabolism impairs chondrocyte function in osteoarthritis,” Osteoarthritis Cartilage, 31(5), pp. 613–626.

[7] Huang, C. Y., Loo, D. M., and Gu, W., 2023, “Modeling of glycosaminoglycan biosynthesis in intervertebral disc cells,” Comput Biol Med, 162, p. 107039.

[8] Myers, E. P., Gallagher, A., Welfley, A., Battles, S., Gregory, A., Brahmachary, P., Greenwood, M., and June, R. K., 2025, “Targeted analysis of chondrocyte central metabolites in response to cyclical compression and shear deformations,” bioRxiv.

[9] Brahmachary, P. P., Welhaven, H. D., and June, R. K., 2023, “Metabolomic Profiling to Understand Chondrocyte Metabolism,” Methods in Molecular Biology, 2598, pp. 141–156.

[10] Jutila, A. A., Zignego, D. L., Schell, W. J., and June, R. K., 2015, “Encapsulation of Chondrocytes in High-Stiffness Agarose Microenvironments for In Vitro Modeling of Osteoarthritis Mechanotransduction,” Annals of Biomedical Engineering, 43(5), pp. 1132–1144.

[11] Romero, P., Wagg, J., Green, M. L., Kaiser, D., Krummenacker, M., and Karp, P. D., 2005, “Computational prediction of human metabolic pathways from the complete human genome,” Genome biology, 6(1), pp. 1–17.

[12] Romero, P., Wagg, J., Green, M. L., Kaiser, D., Krummenacker, M., and Karp, P. D., 2005, “Computational prediction of human metabolic pathways from the complete human genome,” Genome Biol, 6(1), p. R2.

[13] UniProt, C., 2023, “UniProt: the Universal Protein Knowledgebase in 2023,” Nucleic Acids Res, 51(D1), pp. D523–D531.

[14] von Kamp, A., Thiele, S., Hadicke, O., and Klamt, S., 2017, “Use of CellNetAnalyzer in biotechnology and metabolic engineering,” J Biotechnol, 261, pp. 221–228.

[15] Chong, P. P., Panjavarnam, P., Ahmad, W., Chan, C. K., Abbas, A. A., Merican, A. M., Pingguan-Murphy, B., and Kamarul, T., 2020, “Mechanical compression controls the biosynthesis of human osteoarthritic chondrocytes in vitro,” Clin Biomech (Bristol), 79, p. 105178.

[16] Takemoto, M., Sugishita, Y., Takahashi-Suzuki, Y., Fujiya, H., Niki, H., and Yudoh, K., 2025, “Repetitive Compressive Loading Downregulates Mitochondria Function and Upregulates the Cartilage Matrix Degrading Enzyme MMP-13 Through the Coactivation of NAD-Dependent Sirtuin 1 and Runx2 in Osteoarthritic Chondrocytes,” Int J Mol Sci, 26(11).

[17] Lane Smith, R., Trindade, M. C., Ikenoue, T., Mohtai, M., Das, P., Carter, D. R., Goodman, S. B., and Schurman, D. J., 2000, “Effects of shear stress on articular chondrocyte metabolism,” Biorheology, 37(1-2), pp. 95–107.

[18] Salinas, D., Minor, C. A., Carlson, R. P., McCutchen, C. N., Mumey, B. M., and June, R. K., 2017, “Combining Targeted Metabolomic Data with a Model of Glucose Metabolism: Toward Progress in Chondrocyte Mechanotransduction,” PloS one, 12(1).

