## Supplemental Material for "Expanded stoichiometric model of chondrocyte metabolism: response to cyclical shear and compressive loading"

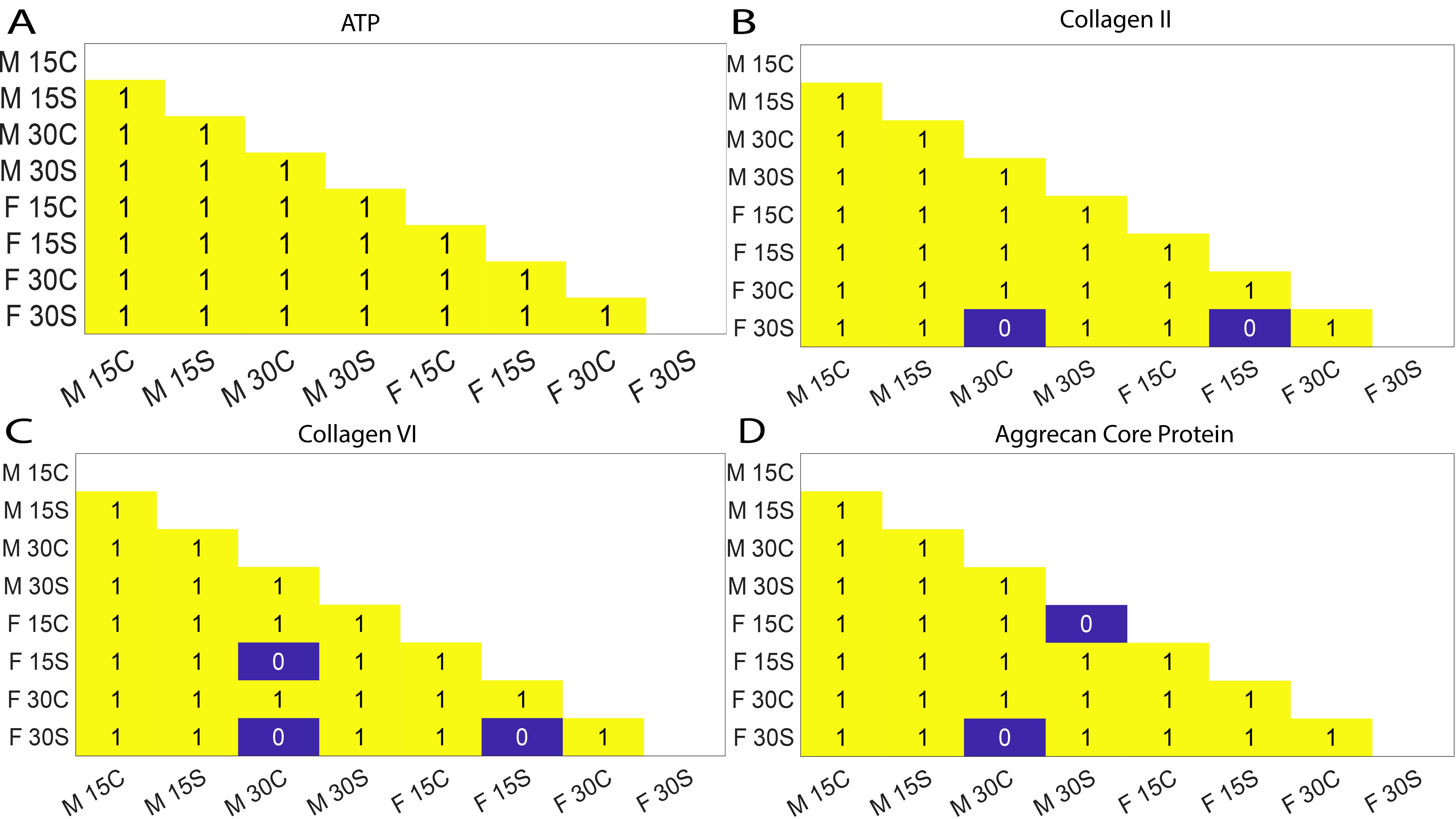

**Figure S1**: Significance tests of predicted flux distributions between experimental groups for maximizers of (a) ATP, (b) collagen II production, (c) collagen VI production, and (d) aggrecan core protein production.. Each cell with a 1 (yellow) indicates that the two compared group’s maximizer distributions are statistically different when comparing the row group versus the column group. Group comparisons with a 0 (purple cells) in the cell represent distributions that were not statistically different from each other when comparing the row group versus the column group. Statistical significance determined via the Kolmogorov-Smirnoff test with Bonferroni-adjusted p-values. This suggests that for the 1000 simulations where ATP was maximized, the distributions of ATP production for each group were statistically different from every other group as seen in S6A. We also see that for collagen II, collagen VI, and aggrecan core protein, that the Male 30C group and F 30S group distributions of production for each of the three maximizers were not different from each other statistically, shown in S6B-D. For the simulations where collagen II was maximized, in addition to the Male 30C group being not statistically different from the Female 30S group, we also see that the Female 15S and Female 30S groups were not statistically different from each other either. The rest of the group comparisons for collagen II show that they were statistically different from each other. For the simulations where collagen VI was maximized, in addition to the Male 30C group being not statistically different from the Female 30S group and the Female 15S and Female 30S groups, like for collagen II, we also see that the Male 30C group and Female 15S group were not statistically different from each other. The rest of the group comparisons show that they were statistically different from each other. For the simulations where aggrecan core protein was maximized, in addition to the Male 30C group not being statistically different from the Female 30S group, like with collagen II and VI, we see that the Male 30C and Female 15S group were also not statistically different from each other. The rest of the groups for aggrecan core protein were statistically different from each other.

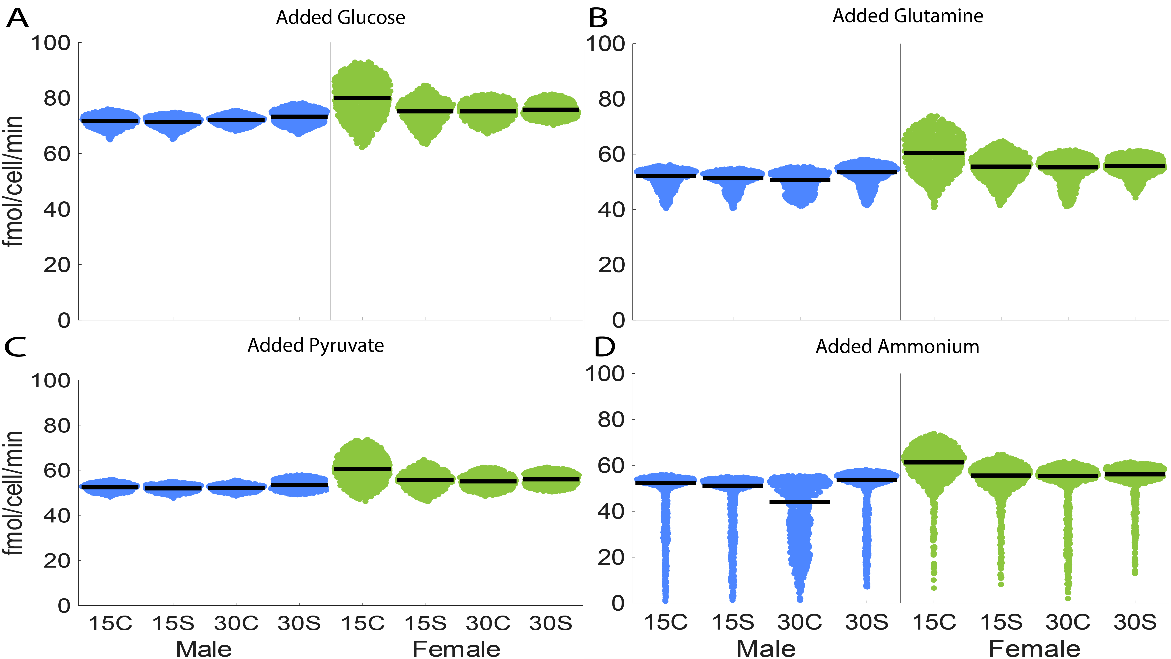

**Figure S2:** Distributions of each group's production of ATP where (A) glucose uptake, (B) glutamine uptake, (C) pyruvate uptake, and (D) ammonium uptake was allowed up to 10 fmol/cell/min in addition to the observed metabolite values as sources. 1000 simulations were run for each group. Simulation results for the male samples in blue and female samples in green. Each experimental condition is represented by a separate distribution from left to right as 15 minutes of compression (15C), 15 minutes of shear (15S), 30 minutes of compression (30C), and 30 minutes of shear (30S). The median maximum flux is denoted by the black bar in each plot. Each point in each distribution represents the maximum ATP flux (in fmol/cell/min) from each simulation.

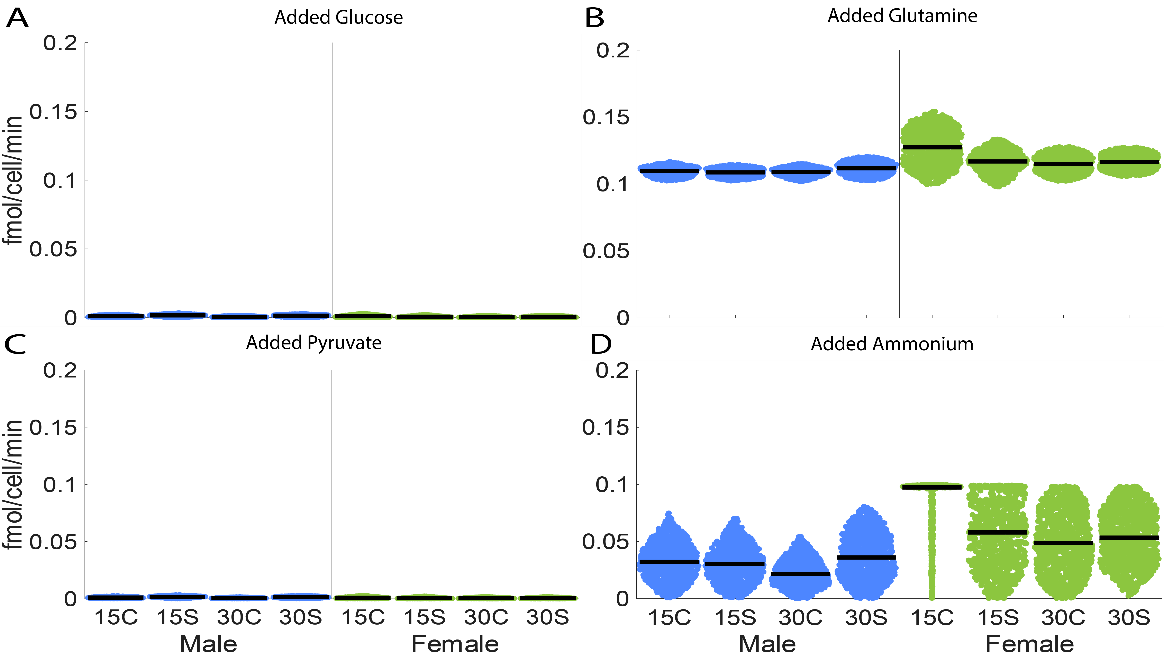

**Figure S3:** Distributions of each group's production of type II collagen where (A) glucose uptake, (B) glutamine uptake, (C) pyruvate uptake, and (D) ammonium uptake was allowed up to 10 fmol/cell/min in addition to the observed metabolite values as sources. 1000 simulations were run for each group. Simulation results for the male samples are in blue and for the female samples are in green. Each experimental condition is represented by a separate distribution from left to right as 15 minutes of compression (15C), 15 minutes of shear (15S), 30 minutes of compression (30C), and 30 minutes of shear (30S). The median maximum flux is denoted by the black bar in each plot. Each point in each distribution represents the maximum type II collagen flux (in fmol/cell/min) from each simulation. This figure suggests that type II collagen production is increased the most by allowing glutamine uptake in our model. The glutamine uptake likely results in a larger increase since it is both a carbon and a nitrogen source, whereas ammonium is only a nitrogen source. This suggests that our system is likely nitrogen-limited, but not likely carbon limited, as collagen II production does not appear to improve with the allowed uptake of glucose or pyruvate.

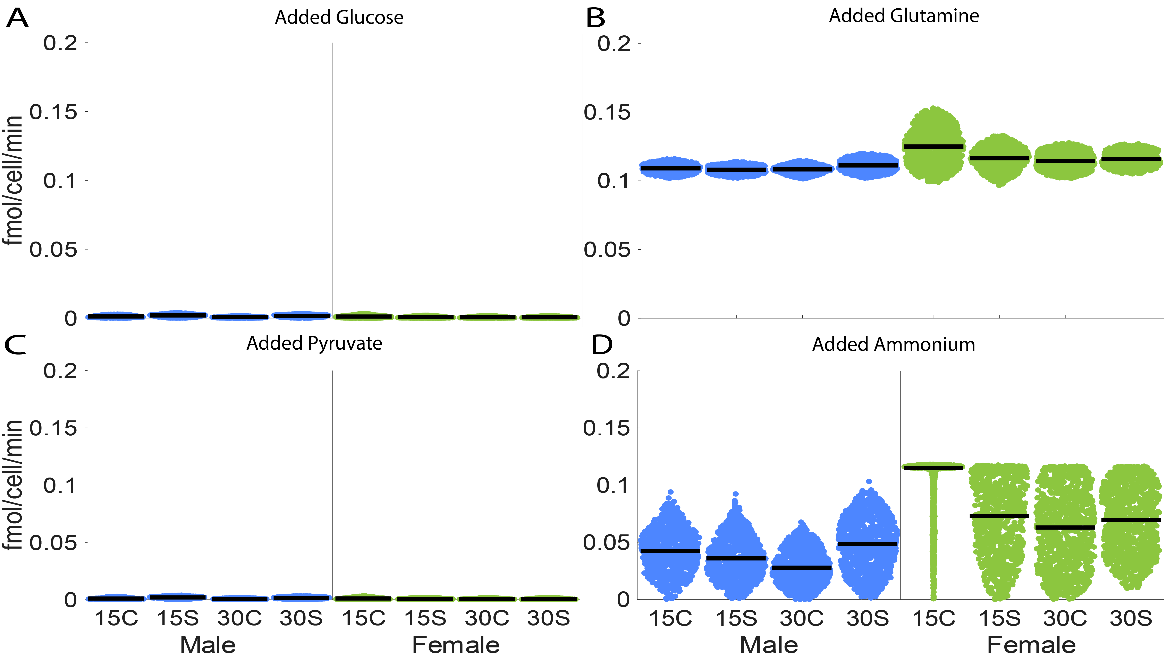

**Figure S4:** Distributions of each group's production of type VI collagen where (A) glucose uptake, (B) glutamine uptake, (C) pyruvate uptake, and (D) ammonium uptake was allowed up to 10 fmol/cell/min in addition to the observed metabolite data as sources. 1000 simulations were run for each group. Simulation results for the male samples are in blue and for the female samples are in green. Each experimental condition is represented by a separate distribution from left to right as 15 minutes of compression (15C), 15 minutes of shear (15S), 30 minutes of compression (30C), and 30 minutes of shear (30S). The median maximum flux is denoted by the black bar in each plot. Each point in each distribution represents the maximum type VI collagen flux (in fmol/cell/min) from each simulation. Similar to collagen II, this figure suggests that collagen VI production is increased the most by allowing glutamine uptake in our model, as seen in S4B. Collagen VI production also increases by allowing ammonium uptake, as seen in S4D. The glutamine uptake likely results in a larger increase since it is both a carbon and a nitrogen source, whereas ammonium is only a nitrogen source. This suggests that our model is likely nitrogen-limited, but not likely carbon limited, as type VI collagen production does not appear to improve with the allowed uptake of glucose or pyruvate.

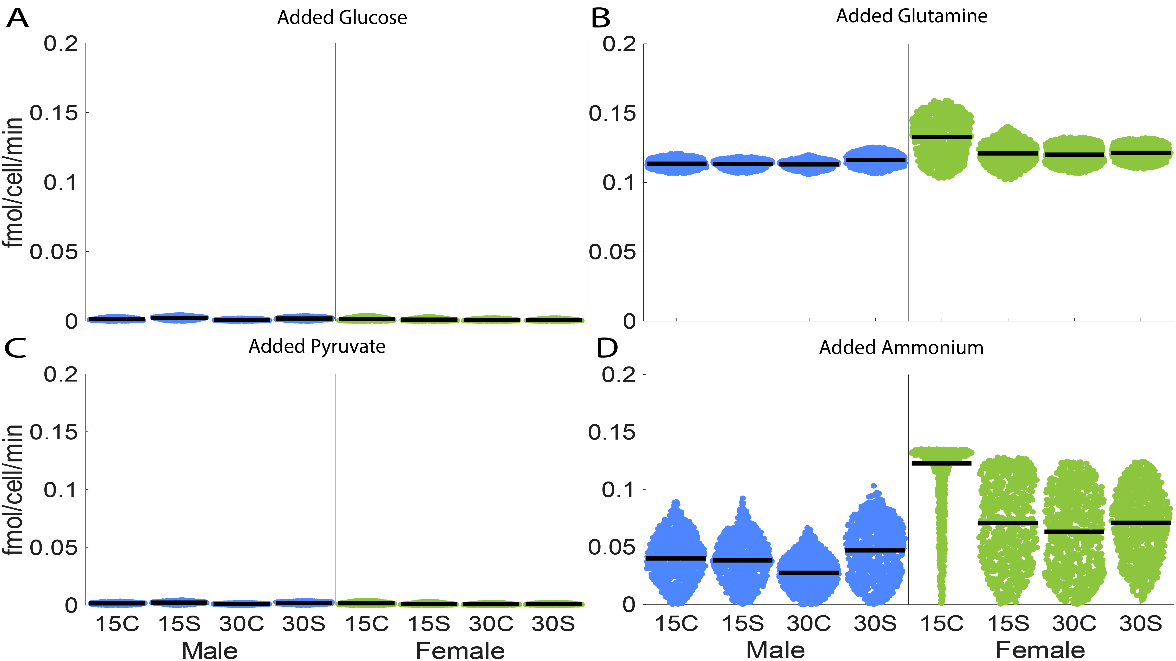

**Figure S5:** Distributions of each group's production of aggrecan where (A) glucose uptake, (B) glutamine uptake, (C) pyruvate uptake, and (D) ammonium uptake was allowed up to 10 fmol/cell/min in addition to the observed metabolite data as sources. 1000 simulations were run for each group. Simulation results for the male samples are in blue and for the female samples are in green. Each experimental condition is represented by a separate distribution from left to right as 15 minutes of compression (15C), 15 minutes of shear (15S), 30 minutes of compression (30C), and 30 minutes of shear (30S). The median maximum flux is denoted by the black bar in each plot. Each point in each distribution represents the maximum aggrecan core protein flux (in fmol/cell/min) from each simulation. Similar to both collagens, this figure suggests that aggrecan production is increased the most by allowing glutamine uptake in our model, as seen in S5B. Aggrecan production also increases by allowing ammonium uptake, as seen in S5D. The glutamine uptake likely results in a larger increase since it is both a carbon and a nitrogen source, whereas ammonium is only a nitrogen source. This suggests that our model is likely nitrogen-limited, but not likely carbon limited, as aggrecan core protein production does not appear to improve with the allowed uptake of glucose or pyruvate.

**Supplemental Tables:**

**Table S1: Maximum possible** ATP and Lactate Yields (mol/mol) on Base Model (no observed metabolite sources) in aerobic and anaerobic conditions. ATP and lactate yields match theoretical values (Lehninger, Nelson, and Cox 2008. Lehninger Principles of Biochemistry), confirming model functionality.

|  | **ATP Yield on Glucose** | **ATP Yield on Pyruvate** | **ATP Yield on Glutamine** | **Lactate Yield on Glucose** | **Lactate Yield on Glutamine** |
| --- | --- | --- | --- | --- | --- |
| **Aerobic** | 32 | 12.5 | 16.525 | NA | NA |
| **Anaerobic** | 2 | None | None | -2 | None |

**Table S2:** ATP production as a function of O_2_ availability, to confirm model functionality. Five separate simulations were run with no measured metabolite data, glucose uptake allowed up to 10 fmol/cell/min, and maximum O_2_uptake allowance decreasing for each simulation on an evenly spaced interval from 10 to 0 fmol/cell/min. As expected, ATP production decreases as oxygen uptake is limited.

|  | **O2 = 10** | **O2 = 7.5** | **O2 = 5** | **O2 = 2.5** | **O2 = 0** |
| --- | --- | --- | --- | --- | --- |
| **ATP** | 70 | 57.5 | 45 | 32.5 | 20 |
