## Supplementary material for "Expanded stoichiometric model of chondrocyte metabolism: response to cyclical shear and compressive loading": SI_2.m

```
%% Ebru chondrocyte model, supplemental file for analysis in CobraToolbox
% Analysis begins with confirmation of basic model properties limodel_27july2023.xmlke ATPchondrocyte_model_27july2023_v4_Ammonium_Source
% yield, collagen yield on glucose and glutamine. This analysis will
% demonstrate that the model is a realistic portrayal of chondrocyte
% metabolism. Note that glutamine is a nonessential amino acid but is
% present in the media that the experiments were run in.
%
% Then the analysis of cost of synthesis of individual nonessential amino acids and
% cost of transport of individual essential amino acids
% 
% Comparison of yields and amino acid costs with high and low O2
% %
% note that the dashes in the CNA model get turned into underscores
%%
initCobraToolbox()
%% Set some reaction names and variables
model = readCbModel('replace_with_xml_file_path','fileType','SBML')

% biomass and ATP maintenance terms
collagenII = find(strcmp('T_Collagen_II',model.rxns));
collagenVI = find(strcmp('T_Collagen_VI',model.rxns));
O2 = find(strcmp('T_O2',model.rxns));
CO2 = find(strcmp('T_CO2',model.rxns));
HCO3 = find(strcmp('T_HCO3',model.rxns));
ATPm = find(strcmp('T_ATP',model.rxns));
glucose_noCost = find(strcmp('T_Glucose_1',model.rxns));
glucose_Na = find(strcmp('T_Glucose_2',model.rxns));
glutamine_noCost = find(strcmp('T_Glutamine_1',model.rxns));
glutamine_Na = find(strcmp('T_Glutamine_2',model.rxns));

alanine = find(strcmp('cost_alanine',model.rxns));
arginine = find(strcmp('cost_arginine',model.rxns));
asparagine = find(strcmp('cost_asparagine',model.rxns));
aspartate = find(strcmp('cost_aspartate',model.rxns));
cysteine = find(strcmp('cost_cysteine',model.rxns));
glutamate = find(strcmp('cost_glutamate',model.rxns));
glutamine = find(strcmp('cost_glutamine',model.rxns));
glycine = find(strcmp('cost_glycine',model.rxns));
proline = find(strcmp('cost_proline',model.rxns));
serine = find(strcmp('cost_serine',model.rxns));
tyrosine = find(strcmp('cost_tyrosine',model.rxns));
phenylalanine = find(strcmp('T_Phe',model.rxns));

% carbon content of each metabolite of interest
Cmol_collagenII = 395;
Cmol_collagenVI = 462;
Cmol_glucose = 6;
Cmol_glutamine = 5;

Cmol_alanine = 3;
Cmol_arginine = 6;
Cmol_asparagine = 4;
Cmol_aspartate = 4;
Cmol_cysteine = 3;
Cmol_glutamate = 5;
Cmol_glycine = 2;
Cmol_proline = 5;
Cmol_serine = 3;
Cmol_tyrosine = 9;
Cmol_phenylalanine = 9;

%% Model predictions for ATPm production on glucose
% first, start with aerobic conditions, use the glucose transporter that
% requires Na+
model = readCbModel('replace_with_xml_file_path','fileType','SBML')
% % prevent the export of nonessential amino acids; aren't needed for this
model = changeRxnBounds(model,'cost_alanine',0,'b');
model = changeRxnBounds(model,'cost_arginine',0,'b');
model = changeRxnBounds(model,'cost_asparagine',0,'b');
model = changeRxnBounds(model,'cost_aspartate',0,'b');
model = changeRxnBounds(model,'cost_cysteine',0,'b');
model = changeRxnBounds(model,'cost_glutamate',0,'b');
model = changeRxnBounds(model,'cost_glutamine',0,'b');
model = changeRxnBounds(model,'cost_glycine',0,'b');
model = changeRxnBounds(model,'cost_proline',0,'b');
model = changeRxnBounds(model,'cost_serine',0,'b');
model = changeRxnBounds(model,'cost_tyrosine',0,'b');
% turn offGTPm
model = changeRxnBounds(model,'T_GTP',0,'b');
% turn off pyruvate uptake
model = changeRxnBounds(model,'T_Pyruvate',0,'u');
% turn off lactate uptake
model = changeRxnBounds(model,'T_Lactate',0,'l');
% turn off CO2 uptake
model = changeRxnBounds(model,'T_CO2',0,'l');
% turn off HCO3 uptake
model = changeRxnBounds(model,'T_HCO3',1,'u');
% have to force transhydrogenase in the forward direction for the
% predictions to run properly. Otherwise, there seems to be an issue with
% Cobratoolbox predictions and ATP production is not actually maximized
model = changeRxnBounds(model,'TransHyd',0,'b');
% prevent glutamine uptake
model = changeRxnBounds(model,'T_Glutamine_1',0,'b');
model = changeRxnBounds(model,'T_Glutamine_2',0,'b');
% turn off glucose transporter that does not require sodium
model = changeRxnBounds(model,'T_Glucose_1',0,'b');
% prevent synthesis of collagen VI and collagen II
model = changeRxnBounds(model,'CollagenVI',0,'b');
model = changeRxnBounds(model,'CollagenII',0,'b');
% set maximization criteria to ATPm
model = changeObjective(model,'T_ATP');
% maximize production of ATP
FBAsolution1 = optimizeCbModel(model,'max');
% pull the important values
yield_ATPm_glucoseNa =((FBAsolution1.v(ATPm))./(FBAsolution1.v(glucose_Na)));

% aerobic conditions but use glucose transporter with no cost
model = readCbModel('replace_with_xml_file_path','fileType','SBML')
% % prevent the export of nonessential amino acids; aren't needed for this
model = changeRxnBounds(model,'cost_alanine',0,'b');
model = changeRxnBounds(model,'cost_arginine',0,'b');
model = changeRxnBounds(model,'cost_asparagine',0,'b');
model = changeRxnBounds(model,'cost_aspartate',0,'b');
model = changeRxnBounds(model,'cost_cysteine',0,'b');
model = changeRxnBounds(model,'cost_glutamate',0,'b');
model = changeRxnBounds(model,'cost_glutamine',0,'b');
model = changeRxnBounds(model,'cost_glycine',0,'b');
model = changeRxnBounds(model,'cost_proline',0,'b');
model = changeRxnBounds(model,'cost_serine',0,'b');
model = changeRxnBounds(model,'cost_tyrosine',0,'b');
% turn offGTPm
model = changeRxnBounds(model,'T_GTP',0,'b');
% turn off pyruvate uptake
model = changeRxnBounds(model,'T_Pyruvate',0,'u');
% turn off lactate uptake
model = changeRxnBounds(model,'T_Lactate',0,'l');
% turn off CO2 uptake
model = changeRxnBounds(model,'T_CO2',0,'l');
% turn off HCO3 uptake
model = changeRxnBounds(model,'T_HCO3',1,'u');
% have to force transhydrogenase in the forward direction for the
% predictions to run properly. Otherwise, there seems to be an issue with
% Cobratoolbox predictions and ATP production is not actually maximized
model = changeRxnBounds(model,'TransHyd',0,'b');
% prevent glutamine uptake
model = changeRxnBounds(model,'T_Glutamine_1',0,'b');
model = changeRxnBounds(model,'T_Glutamine_2',0,'b');
% turn off glucose transporter that requires sodium
model = changeRxnBounds(model,'T_Glucose_2',0,'b');
% prevent synthesis of collagen VI and collagen II
model = changeRxnBounds(model,'CollagenVI',0,'b');
model = changeRxnBounds(model,'CollagenII',0,'b');
% set maximization criteria to ATPm
model = changeObjective(model,'T_ATP');
% checking transhydrogenase reaction
model = changeRxnBounds(model,'TransHyd',0,'l');
% maximize production of collagen
FBAsolution2 = optimizeCbModel(model,'max');
% pull the important values
yield_ATPm_glucosenoCost =((FBAsolution2.v(ATPm))./(FBAsolution2.v(glucose_noCost)));

% prevent use of O2 as terminal electron acceptor, use glucose transporter that requires Na+
model = readCbModel('replace_with_xml_file_path','fileType','SBML')
% % prevent the export of nonessential amino acids; aren't needed for this
model = changeRxnBounds(model,'cost_alanine',0,'b');
model = changeRxnBounds(model,'cost_arginine',0,'b');
model = changeRxnBounds(model,'cost_asparagine',0,'b');
model = changeRxnBounds(model,'cost_aspartate',0,'b');
model = changeRxnBounds(model,'cost_cysteine',0,'b');
model = changeRxnBounds(model,'cost_glutamate',0,'b');
model = changeRxnBounds(model,'cost_glutamine',0,'b');
model = changeRxnBounds(model,'cost_glycine',0,'b');
model = changeRxnBounds(model,'cost_proline',0,'b');
model = changeRxnBounds(model,'cost_serine',0,'b');
model = changeRxnBounds(model,'cost_tyrosine',0,'b');
% turn offGTPm
model = changeRxnBounds(model,'T_GTP',0,'b');
% turn off pyruvate uptake
model = changeRxnBounds(model,'T_Pyruvate',0,'u');
% turn off lactate uptake
model = changeRxnBounds(model,'T_Lactate',0,'l');
% turn off CO2 uptake
model = changeRxnBounds(model,'T_CO2',0,'l');
% turn off HCO3 uptake
model = changeRxnBounds(model,'T_HCO3',1,'u');
% have to force transhydrogenase in the forward direction for the
% predictions to run properly. Otherwise, there seems to be an issue with
% Cobratoolbox predictions and ATP production is not actually maximized
model = changeRxnBounds(model,'TransHyd',0,'b');
% prevent glutamine uptake
model = changeRxnBounds(model,'T_Glutamine_1',0,'b');
model = changeRxnBounds(model,'T_Glutamine_2',0,'b');
% turn off glucose transporter that does not require sodium
model = changeRxnBounds(model,'T_Glucose_1',0,'b');
% prevent synthesis of collagen VI and collagen II
model = changeRxnBounds(model,'CollagenVI',0,'b');
model = changeRxnBounds(model,'CollagenII',0,'b');
% prevent use of aerobic electron transport chain
model = changeRxnBounds(model,'ETC_4',0,'b');
% set maximization criteria to ATPm
model = changeObjective(model,'T_ATP');
% maximize production of collagen
FBAsolution3 = optimizeCbModel(model,'max');
% pull the important values
yield_ATPm_glucoseNa_noETC4 =((FBAsolution3.v(ATPm))./(FBAsolution3.v(glucose_Na)));

% no O2 as terminal electron acceptor; glucose transporter with no cost
model = readCbModel('replace_with_xml_file_path','fileType','SBML')
% % prevent the export of nonessential amino acids; aren't needed for this
model = changeRxnBounds(model,'cost_alanine',0,'b');
model = changeRxnBounds(model,'cost_arginine',0,'b');
model = changeRxnBounds(model,'cost_asparagine',0,'b');
model = changeRxnBounds(model,'cost_aspartate',0,'b');
model = changeRxnBounds(model,'cost_cysteine',0,'b');
model = changeRxnBounds(model,'cost_glutamate',0,'b');
model = changeRxnBounds(model,'cost_glutamine',0,'b');
model = changeRxnBounds(model,'cost_glycine',0,'b');
model = changeRxnBounds(model,'cost_proline',0,'b');
model = changeRxnBounds(model,'cost_serine',0,'b');
model = changeRxnBounds(model,'cost_tyrosine',0,'b');
% turn offGTPm
model = changeRxnBounds(model,'T_GTP',0,'b');
% turn off pyruvate uptake
model = changeRxnBounds(model,'T_Pyruvate',0,'u');
% turn off lactate uptake
model = changeRxnBounds(model,'T_Lactate',0,'l');
% turn off CO2 uptake
model = changeRxnBounds(model,'T_CO2',0,'l');
% turn off HCO3 uptake
model = changeRxnBounds(model,'T_HCO3',1,'u');
% have to force transhydrogenase in the forward direction for the
% predictions to run properly. Otherwise, there seems to be an issue with
% Cobratoolbox predictions and ATP production is not actually maximized
model = changeRxnBounds(model,'TransHyd',0,'b');
% prevent glutamine uptake
model = changeRxnBounds(model,'T_Glutamine_1',0,'b');
model = changeRxnBounds(model,'T_Glutamine_2',0,'b');
% turn off glucose transporter that requires sodium
model = changeRxnBounds(model,'T_Glucose_2',0,'b');
% prevent use of O2 as terminal electron acceptor
model = changeRxnBounds(model,'ETC_4',0,'b');
% prevent synthesis of collagen VI and collagen II
model = changeRxnBounds(model,'CollagenVI',0,'b');
model = changeRxnBounds(model,'CollagenII',0,'b');
% set maximization criteria to ATPm
model = changeObjective(model,'T_ATP');
% maximize production of collagen
FBAsolution4 = optimizeCbModel(model,'max');
% pull the important values
yield_ATPm_glucosenoCost_noETC4 =((FBAsolution4.v(ATPm))./(FBAsolution4.v(glucose_noCost)));

%% Model predictions for collagen II yields on glucose 

% first, start with aerobic conditions, use the glucose transporter that
% requires Na+
model = readCbModel('replace_with_xml_file_path','fileType','SBML')
% % prevent the export of nonessential amino acids; aren't needed for this
model = changeRxnBounds(model,'cost_alanine',0,'b');
model = changeRxnBounds(model,'cost_arginine',0,'b');
model = changeRxnBounds(model,'cost_asparagine',0,'b');
model = changeRxnBounds(model,'cost_aspartate',0,'b');
model = changeRxnBounds(model,'cost_cysteine',0,'b');
model = changeRxnBounds(model,'cost_glutamate',0,'b');
model = changeRxnBounds(model,'cost_glutamine',0,'b');
model = changeRxnBounds(model,'cost_glycine',0,'b');
model = changeRxnBounds(model,'cost_proline',0,'b');
model = changeRxnBounds(model,'cost_serine',0,'b');
model = changeRxnBounds(model,'cost_tyrosine',0,'b');
% turn off ATPm and GTPm
model = changeRxnBounds(model,'T_ATP',0,'b');
model = changeRxnBounds(model,'T_GTP',0,'b');
% turn off pyruvate uptake
model = changeRxnBounds(model,'T_Pyruvate',0,'u');
% turn off lactate uptake
model = changeRxnBounds(model,'T_Lactate',0,'l');
% turn off CO2 uptake
model = changeRxnBounds(model,'T_CO2',0,'l');
% turn off HCO3 uptake
model = changeRxnBounds(model,'T_HCO3',1,'u');
% have to force transhydrogenase in the forward direction for the
% predictions to run properly. Otherwise, there seems to be an issue with
% Cobratoolbox predictions and ATP production is not actually maximized
model = changeRxnBounds(model,'TransHyd',0,'b');
% prevent glutamine uptake
model = changeRxnBounds(model,'T_Glutamine_1',0,'b');
model = changeRxnBounds(model,'T_Glutamine_2',0,'b');
% turn off glucose transporter that does not require sodium
model = changeRxnBounds(model,'T_Glucose_1',0,'b');
% prevent synthesis of collagen VI
model = changeRxnBounds(model,'CollagenVI',0,'b');
% set maximization criteria to collagen II
model = changeObjective(model,'T_Collagen_II');
% maximize production of collagen
FBAsolution1 = optimizeCbModel(model,'max');
% pull the important values
yield_CollagenII_glucoseNa =((FBAsolution1.v(collagenII)*Cmol_collagenII)./(FBAsolution1.v(glucose_Na)*Cmol_glucose));

% next, aerobic conditions, use the transporter with no cost
model = readCbModel('replace_with_xml_file_path','fileType','SBML')
% prevent the export of nonessential amino acids; aren't needed for this
model = changeRxnBounds(model,'cost_alanine',0,'b');
model = changeRxnBounds(model,'cost_arginine',0,'b');
model = changeRxnBounds(model,'cost_asparagine',0,'b');
model = changeRxnBounds(model,'cost_aspartate',0,'b');
model = changeRxnBounds(model,'cost_cysteine',0,'b');
model = changeRxnBounds(model,'cost_glutamate',0,'b');
model = changeRxnBounds(model,'cost_glutamine',0,'b');
model = changeRxnBounds(model,'cost_glycine',0,'b');
model = changeRxnBounds(model,'cost_proline',0,'b');
model = changeRxnBounds(model,'cost_serine',0,'b');
model = changeRxnBounds(model,'cost_tyrosine',0,'b');
% turn off ATPm and GTPm
model = changeRxnBounds(model,'T_ATP',0,'b');
model = changeRxnBounds(model,'T_GTP',0,'b');
% turn off pyruvate uptake
model = changeRxnBounds(model,'T_Pyruvate',0,'u');
% turn off lactate uptake
model = changeRxnBounds(model,'T_Lactate',0,'l');
% turn off CO2 uptake
model = changeRxnBounds(model,'T_CO2',0,'l');
% turn off HCO3 uptake
model = changeRxnBounds(model,'T_HCO3',1,'u');
% have to force transhydrogenase in the forward direction for the
% predictions to run properly. Otherwise, there seems to be an issue with
% Cobratoolbox predictions and ATP production is not actually maximized
model = changeRxnBounds(model,'TransHyd',0,'b');
% prevent glutamine uptake
model = changeRxnBounds(model,'T_Glutamine_1',0,'b');
model = changeRxnBounds(model,'T_Glutamine_2',0,'b');
% turn off glucose transporter that requires sodium
model = changeRxnBounds(model,'T_Glucose_2',0,'b');
% prevent synthesis of collagen VI
model = changeRxnBounds(model,'CollagenVI',0,'b');
% set maximization criteria to collagen II
model = changeObjective(model,'T_Collagen_II');
% maximize production of collagen
FBAsolution2 = optimizeCbModel(model,'max');
% pull the important values
yield_CollagenII_glucosenoCost =((FBAsolution2.v(collagenII)*Cmol_collagenII)./(FBAsolution2.v(glucose_noCost)*Cmol_glucose));

% no O2 as terminal electron acceptor, use the glucose transporter that requires Na+
model = readCbModel('replace_with_xml_file_path','fileType','SBML')
% prevent the export of nonessential amino acids; aren't needed for this
model = changeRxnBounds(model,'cost_alanine',0,'b');
model = changeRxnBounds(model,'cost_arginine',0,'b');
model = changeRxnBounds(model,'cost_asparagine',0,'b');
model = changeRxnBounds(model,'cost_aspartate',0,'b');
model = changeRxnBounds(model,'cost_cysteine',0,'b');
model = changeRxnBounds(model,'cost_glutamate',0,'b');
model = changeRxnBounds(model,'cost_glutamine',0,'b');
model = changeRxnBounds(model,'cost_glycine',0,'b');
model = changeRxnBounds(model,'cost_proline',0,'b');
model = changeRxnBounds(model,'cost_serine',0,'b');
model = changeRxnBounds(model,'cost_tyrosine',0,'b');
% turn off ATPm and GTPm
model = changeRxnBounds(model,'T_ATP',0,'b');
model = changeRxnBounds(model,'T_GTP',0,'b');
% turn off pyruvate uptake
model = changeRxnBounds(model,'T_Pyruvate',0,'u');
% turn off lactate uptake
model = changeRxnBounds(model,'T_Lactate',0,'l');
% turn off CO2 uptake
model = changeRxnBounds(model,'T_CO2',0,'l');
% turn off HCO3 uptake
model = changeRxnBounds(model,'T_HCO3',1,'u');
% have to force transhydrogenase in the forward direction for the
% predictions to run properly. Otherwise, there seems to be an issue with
% Cobratoolbox predictions and ATP production is not actually maximized
model = changeRxnBounds(model,'TransHyd',0,'b');
% prevent glutamine uptake
model = changeRxnBounds(model,'T_Glutamine_1',0,'b');
model = changeRxnBounds(model,'T_Glutamine_2',0,'b');
% turn off glucose transporter that does not require sodium
model = changeRxnBounds(model,'T_Glucose_1',0,'b');
% prevent synthesis of collagen VI
model = changeRxnBounds(model,'CollagenVI',0,'b');
% prevent use of O2 as terminal electron acceptor
model = changeRxnBounds(model,'ETC_4',0,'b');
% set maximization criteria to collagen II
model = changeObjective(model,'T_Collagen_II');
% maximize production of collagen
FBAsolution3 = optimizeCbModel(model,'max');
% pull the important values
yield_CollagenII_glucoseNa_noETC4 =((FBAsolution3.v(collagenII)*Cmol_collagenII)./(FBAsolution3.v(glucose_Na)*Cmol_glucose));

% no O2 as terminal electron acceptor, use the no cost glucose transporter
model = readCbModel('replace_with_xml_file_path','fileType','SBML')
% prevent the export of nonessential amino acids; aren't needed for this
model = changeRxnBounds(model,'cost_alanine',0,'b');
model = changeRxnBounds(model,'cost_arginine',0,'b');
model = changeRxnBounds(model,'cost_asparagine',0,'b');
model = changeRxnBounds(model,'cost_aspartate',0,'b');
model = changeRxnBounds(model,'cost_cysteine',0,'b');
model = changeRxnBounds(model,'cost_glutamate',0,'b');
model = changeRxnBounds(model,'cost_glutamine',0,'b');
model = changeRxnBounds(model,'cost_glycine',0,'b');
model = changeRxnBounds(model,'cost_proline',0,'b');
model = changeRxnBounds(model,'cost_serine',0,'b');
model = changeRxnBounds(model,'cost_tyrosine',0,'b');
% turn off ATPm and GTPm
model = changeRxnBounds(model,'T_ATP',0,'b');
model = changeRxnBounds(model,'T_GTP',0,'b');
% turn off pyruvate uptake
model = changeRxnBounds(model,'T_Pyruvate',0,'u');
% turn off lactate uptake
model = changeRxnBounds(model,'T_Lactate',0,'l');
% turn off CO2 uptake
model = changeRxnBounds(model,'T_CO2',0,'l');
% turn off HCO3 uptake
model = changeRxnBounds(model,'T_HCO3',1,'u');
% have to force transhydrogenase in the forward direction for the
% predictions to run properly. Otherwise, there seems to be an issue with
% Cobratoolbox predictions and ATP production is not actually maximized
model = changeRxnBounds(model,'TransHyd',0,'b');
% prevent glutamine uptake
model = changeRxnBounds(model,'T_Glutamine_1',0,'b');
model = changeRxnBounds(model,'T_Glutamine_2',0,'b');
% turn off glucose transporter that does not require sodium
model = changeRxnBounds(model,'T_Glucose_2',0,'b');
% prevent synthesis of collagen VI
model = changeRxnBounds(model,'CollagenVI',0,'b');
% prevent use of O2 as terminal electron acceptor
model = changeRxnBounds(model,'ETC_4',0,'b');
% set maximization criteria to collagen II
model = changeObjective(model,'T_Collagen_II');
% maximize production of collagen
FBAsolution4 = optimizeCbModel(model,'max');
% pull the important values
yield_CollagenII_glucosenoCost_noETC4 =((FBAsolution4.v(collagenII)*Cmol_collagenII)./(FBAsolution4.v(glucose_noCost)*Cmol_glucose));

%% Model predictions for collagen VI yields on glucose 

% first, start with aerobic conditions, use the glucose transporter that
% requires Na+
model = readCbModel('replace_with_xml_file_path','fileType','SBML')
% % prevent the export of nonessential amino acids; aren't needed for this
model = changeRxnBounds(model,'cost_alanine',0,'b');
model = changeRxnBounds(model,'cost_arginine',0,'b');
model = changeRxnBounds(model,'cost_asparagine',0,'b');
model = changeRxnBounds(model,'cost_aspartate',0,'b');
model = changeRxnBounds(model,'cost_cysteine',0,'b');
model = changeRxnBounds(model,'cost_glutamate',0,'b');
model = changeRxnBounds(model,'cost_glutamine',0,'b');
model = changeRxnBounds(model,'cost_glycine',0,'b');
model = changeRxnBounds(model,'cost_proline',0,'b');
model = changeRxnBounds(model,'cost_serine',0,'b');
model = changeRxnBounds(model,'cost_tyrosine',0,'b');
% turn off ATPm and GTPm
model = changeRxnBounds(model,'T_ATP',0,'b');
model = changeRxnBounds(model,'T_GTP',0,'b');
% turn off pyruvate uptake
model = changeRxnBounds(model,'T_Pyruvate',0,'u');
% turn off lactate uptake
model = changeRxnBounds(model,'T_Lactate',0,'l');
% turn off CO2 uptake
model = changeRxnBounds(model,'T_CO2',0,'l');
% turn off HCO3 uptake
model = changeRxnBounds(model,'T_HCO3',1,'u');
% have to force transhydrogenase in the forward direction for the
% predictions to run properly. Otherwise, there seems to be an issue with
% Cobratoolbox predictions and ATP production is not actually maximized
model = changeRxnBounds(model,'TransHyd',0,'b');
% prevent glutamine uptake
model = changeRxnBounds(model,'T_Glutamine_1',0,'b');
model = changeRxnBounds(model,'T_Glutamine_2',0,'b');
% turn off glucose transporter that does not require sodium
model = changeRxnBounds(model,'T_Glucose_1',0,'b');
% prevent synthesis of collagen II
model = changeRxnBounds(model,'CollagenII',0,'b');
% set maximization criteria to collagen VI
model = changeObjective(model,'T_Collagen_VI');
% maximize production of collagen
FBAsolution1 = optimizeCbModel(model,'max');
% pull the important values
yield_CollagenVI_glucoseNa =((FBAsolution1.v(collagenVI)*Cmol_collagenVI)./(FBAsolution1.v(glucose_Na)*Cmol_glucose));

% next, aerobic conditions, use the transporter with no cost
model = readCbModel('replace_with_xml_file_path','fileType','SBML')
% prevent the export of nonessential amino acids; aren't needed for this
model = changeRxnBounds(model,'cost_alanine',0,'b');
model = changeRxnBounds(model,'cost_arginine',0,'b');
model = changeRxnBounds(model,'cost_asparagine',0,'b');
model = changeRxnBounds(model,'cost_aspartate',0,'b');
model = changeRxnBounds(model,'cost_cysteine',0,'b');
model = changeRxnBounds(model,'cost_glutamate',0,'b');
model = changeRxnBounds(model,'cost_glutamine',0,'b');
model = changeRxnBounds(model,'cost_glycine',0,'b');
model = changeRxnBounds(model,'cost_proline',0,'b');
model = changeRxnBounds(model,'cost_serine',0,'b');
model = changeRxnBounds(model,'cost_tyrosine',0,'b');
% turn off ATPm and GTPm
model = changeRxnBounds(model,'T_ATP',0,'b');
model = changeRxnBounds(model,'T_GTP',0,'b');
% turn off pyruvate uptake
model = changeRxnBounds(model,'T_Pyruvate',0,'u');
% turn off lactate uptake
model = changeRxnBounds(model,'T_Lactate',0,'l');
% turn off CO2 uptake
model = changeRxnBounds(model,'T_CO2',0,'l');
% turn off HCO3 uptake
model = changeRxnBounds(model,'T_HCO3',1,'u');
% have to force transhydrogenase in the forward direction for the
% predictions to run properly. Otherwise, there seems to be an issue with
% Cobratoolbox predictions and ATP production is not actually maximized
model = changeRxnBounds(model,'TransHyd',0,'b');
% prevent glutamine uptake
model = changeRxnBounds(model,'T_Glutamine_1',0,'b');
model = changeRxnBounds(model,'T_Glutamine_2',0,'b');
% turn off glucose transporter that requires sodium
model = changeRxnBounds(model,'T_Glucose_2',0,'b');
% prevent synthesis of collagen II
model = changeRxnBounds(model,'CollagenII',0,'b');
% set maximization criteria to collagen VI
model = changeObjective(model,'T_Collagen_VI');
% maximize production of collagen
FBAsolution2 = optimizeCbModel(model,'max');
% pull the important values
yield_CollagenVI_glucosenoCost =((FBAsolution2.v(collagenVI)*Cmol_collagenVI)./(FBAsolution2.v(glucose_noCost)*Cmol_glucose));

% no O2 as terminal electron acceptor, use the glucose transporter that requires Na+
model = readCbModel('replace_with_xml_file_path','fileType','SBML')
% prevent the export of nonessential amino acids; aren't needed for this
model = changeRxnBounds(model,'cost_alanine',0,'b');
model = changeRxnBounds(model,'cost_arginine',0,'b');
model = changeRxnBounds(model,'cost_asparagine',0,'b');
model = changeRxnBounds(model,'cost_aspartate',0,'b');
model = changeRxnBounds(model,'cost_cysteine',0,'b');
model = changeRxnBounds(model,'cost_glutamate',0,'b');
model = changeRxnBounds(model,'cost_glutamine',0,'b');
model = changeRxnBounds(model,'cost_glycine',0,'b');
model = changeRxnBounds(model,'cost_proline',0,'b');
model = changeRxnBounds(model,'cost_serine',0,'b');
model = changeRxnBounds(model,'cost_tyrosine',0,'b');
% turn off ATPm and GTPm
model = changeRxnBounds(model,'T_ATP',0,'b');
model = changeRxnBounds(model,'T_GTP',0,'b');
% turn off pyruvate uptake
model = changeRxnBounds(model,'T_Pyruvate',0,'u');
% turn off lactate uptake
model = changeRxnBounds(model,'T_Lactate',0,'l');
% turn off CO2 uptake
model = changeRxnBounds(model,'T_CO2',0,'l');
% turn off HCO3 uptake
model = changeRxnBounds(model,'T_HCO3',1,'u');
% have to force transhydrogenase in the forward direction for the
% predictions to run properly. Otherwise, there seems to be an issue with
% Cobratoolbox predictions and ATP production is not actually maximized
model = changeRxnBounds(model,'TransHyd',0,'b');
% prevent glutamine uptake
model = changeRxnBounds(model,'T_Glutamine_1',0,'b');
model = changeRxnBounds(model,'T_Glutamine_2',0,'b');
% turn off glucose transporter that does not require sodium
model = changeRxnBounds(model,'T_Glucose_1',0,'b');
% prevent synthesis of collagen II
model = changeRxnBounds(model,'CollagenII',0,'b');
% prevent use of O2 as terminal electron acceptor
model = changeRxnBounds(model,'ETC_4',0,'b');
% set maximization criteria to collagen VI
model = changeObjective(model,'T_Collagen_VI');
% maximize production of collagen
FBAsolution3 = optimizeCbModel(model,'max');
% pull the important values
yield_CollagenVI_glucoseNa_noETC4 =((FBAsolution3.v(collagenVI)*Cmol_collagenVI)./(FBAsolution3.v(glucose_Na)*Cmol_glucose));

% no O2 as terminal electron acceptor, use the no cost glucose transporter
model = readCbModel('replace_with_xml_file_path','fileType','SBML')
% prevent the export of nonessential amino acids; aren't needed for this
model = changeRxnBounds(model,'cost_alanine',0,'b');
model = changeRxnBounds(model,'cost_arginine',0,'b');
model = changeRxnBounds(model,'cost_asparagine',0,'b');
model = changeRxnBounds(model,'cost_aspartate',0,'b');
model = changeRxnBounds(model,'cost_cysteine',0,'b');
model = changeRxnBounds(model,'cost_glutamate',0,'b');
model = changeRxnBounds(model,'cost_glutamine',0,'b');
model = changeRxnBounds(model,'cost_glycine',0,'b');
model = changeRxnBounds(model,'cost_proline',0,'b');
model = changeRxnBounds(model,'cost_serine',0,'b');
model = changeRxnBounds(model,'cost_tyrosine',0,'b');
% turn off ATPm and GTPm
model = changeRxnBounds(model,'T_ATP',0,'b');
model = changeRxnBounds(model,'T_GTP',0,'b');
% turn off pyruvate uptake
model = changeRxnBounds(model,'T_Pyruvate',0,'u');
% turn off lactate uptake
model = changeRxnBounds(model,'T_Lactate',0,'l');
% turn off CO2 uptake
model = changeRxnBounds(model,'T_CO2',0,'l');
% turn off HCO3 uptake
model = changeRxnBounds(model,'T_HCO3',1,'u');
% have to force transhydrogenase in the forward direction for the
% predictions to run properly. Otherwise, there seems to be an issue with
% Cobratoolbox predictions and ATP production is not actually maximized
model = changeRxnBounds(model,'TransHyd',0,'b');
% prevent glutamine uptake
model = changeRxnBounds(model,'T_Glutamine_1',0,'b');
model = changeRxnBounds(model,'T_Glutamine_2',0,'b');
% turn off glucose transporter that does not require sodium
model = changeRxnBounds(model,'T_Glucose_2',0,'b');
% prevent synthesis of collagen II
model = changeRxnBounds(model,'CollagenII',0,'b');
% prevent use of O2 as terminal electron acceptor
model = changeRxnBounds(model,'ETC_4',0,'b');
% set maximization criteria to collagen VI
model = changeObjective(model,'T_Collagen_VI');
% maximize production of collagen
FBAsolution4 = optimizeCbModel(model,'max');
% pull the important values
yield_CollagenVI_glucosenoCost_noETC4 =((FBAsolution4.v(collagenVI)*Cmol_collagenVI)./(FBAsolution4.v(glucose_noCost)*Cmol_glucose));

%% Model predictions for ATPm production on glucose
% first, start with aerobic conditions, use the glucose transporter that
% requires Na+
model = readCbModel('replace_with_xml_file_path','fileType','SBML')
% % prevent the export of nonessential amino acids; aren't needed for this
model = changeRxnBounds(model,'cost_alanine',0,'b');
model = changeRxnBounds(model,'cost_arginine',0,'b');
model = changeRxnBounds(model,'cost_asparagine',0,'b');
model = changeRxnBounds(model,'cost_aspartate',0,'b');
model = changeRxnBounds(model,'cost_cysteine',0,'b');
model = changeRxnBounds(model,'cost_glutamate',0,'b');
model = changeRxnBounds(model,'cost_glutamine',0,'b');
model = changeRxnBounds(model,'cost_glycine',0,'b');
model = changeRxnBounds(model,'cost_proline',0,'b');
model = changeRxnBounds(model,'cost_serine',0,'b');
model = changeRxnBounds(model,'cost_tyrosine',0,'b');
% turn offGTPm
model = changeRxnBounds(model,'T_GTP',0,'b');
% turn off pyruvate uptake
model = changeRxnBounds(model,'T_Pyruvate',0,'u');
% turn off lactate uptake
model = changeRxnBounds(model,'T_Lactate',0,'l');
% turn off CO2 uptake
model = changeRxnBounds(model,'T_CO2',0,'l');
% turn off HCO3 uptake
model = changeRxnBounds(model,'T_HCO3',1,'u');
% have to force transhydrogenase in the forward direction for the
% predictions to run properly. Otherwise, there seems to be an issue with
% Cobratoolbox predictions and ATP production is not actually maximized
model = changeRxnBounds(model,'TransHyd',0,'b');
% prevent glucose uptake
model = changeRxnBounds(model,'T_Glucose_1',0,'b');
model = changeRxnBounds(model,'T_Glucose_2',0,'b');
% turn off glutamine transporter that does not require sodium
model = changeRxnBounds(model,'T_Glutamine_1',0,'b');
% prevent synthesis of collagen VI and collagen II
model = changeRxnBounds(model,'CollagenVI',0,'b');
model = changeRxnBounds(model,'CollagenII',0,'b');
% set maximization criteria to ATPm
model = changeObjective(model,'T_ATP');
% maximize production of collagen
FBAsolution1 = optimizeCbModel(model,'max');
% pull the important values
yield_ATPm_glutamineNa =((FBAsolution1.v(ATPm))./(FBAsolution1.v(glutamine_Na)));

% aerobic conditions but use glucose transporter with no cost
model = readCbModel('replace_with_xml_file_path','fileType','SBML')
% % prevent the export of nonessential amino acids; aren't needed for this
model = changeRxnBounds(model,'cost_alanine',0,'b');
model = changeRxnBounds(model,'cost_arginine',0,'b');
model = changeRxnBounds(model,'cost_asparagine',0,'b');
model = changeRxnBounds(model,'cost_aspartate',0,'b');
model = changeRxnBounds(model,'cost_cysteine',0,'b');
model = changeRxnBounds(model,'cost_glutamate',0,'b');
model = changeRxnBounds(model,'cost_glutamine',0,'b');
model = changeRxnBounds(model,'cost_glycine',0,'b');
model = changeRxnBounds(model,'cost_proline',0,'b');
model = changeRxnBounds(model,'cost_serine',0,'b');
model = changeRxnBounds(model,'cost_tyrosine',0,'b');
% turn offGTPm
model = changeRxnBounds(model,'T_GTP',0,'b');
% turn off pyruvate uptake
model = changeRxnBounds(model,'T_Pyruvate',0,'u');
% turn off lactate uptake
model = changeRxnBounds(model,'T_Lactate',0,'l');
% turn off CO2 uptake
model = changeRxnBounds(model,'T_CO2',0,'l');
% turn off HCO3 uptake
model = changeRxnBounds(model,'T_HCO3',1,'u');
% have to force transhydrogenase in the forward direction for the
% predictions to run properly. Otherwise, there seems to be an issue with
% Cobratoolbox predictions and ATP production is not actually maximized
model = changeRxnBounds(model,'TransHyd',0,'b');
% prevent glucose uptake
model = changeRxnBounds(model,'T_Glucose_1',0,'b');
model = changeRxnBounds(model,'T_Glucose_2',0,'b');
% turn off glucose transporter that requires sodium
model = changeRxnBounds(model,'T_Glutamine_2',0,'b');
% prevent synthesis of collagen VI and collagen II
model = changeRxnBounds(model,'CollagenVI',0,'b');
model = changeRxnBounds(model,'CollagenII',0,'b');
% set maximization criteria to ATPm
model = changeObjective(model,'T_ATP');
% checking transhydrogenase reaction
model = changeRxnBounds(model,'TransHyd',0,'l');
% maximize production of collagen
FBAsolution2 = optimizeCbModel(model,'max');
% pull the important values
yield_ATPm_glutaminenoCost =((FBAsolution2.v(ATPm))./(FBAsolution2.v(glutamine_noCost)));

% prevent use of O2 as terminal electron acceptor, use glucose transporter that requires Na+
model = readCbModel('replace_with_xml_file_path','fileType','SBML')
% % prevent the export of nonessential amino acids; aren't needed for this
model = changeRxnBounds(model,'cost_alanine',0,'b');
model = changeRxnBounds(model,'cost_arginine',0,'b');
model = changeRxnBounds(model,'cost_asparagine',0,'b');
model = changeRxnBounds(model,'cost_aspartate',0,'b');
model = changeRxnBounds(model,'cost_cysteine',0,'b');
model = changeRxnBounds(model,'cost_glutamate',0,'b');
model = changeRxnBounds(model,'cost_glutamine',0,'b');
model = changeRxnBounds(model,'cost_glycine',0,'b');
model = changeRxnBounds(model,'cost_proline',0,'b');
model = changeRxnBounds(model,'cost_serine',0,'b');
model = changeRxnBounds(model,'cost_tyrosine',0,'b');
% turn offGTPm
model = changeRxnBounds(model,'T_GTP',0,'b');
% turn off pyruvate uptake
model = changeRxnBounds(model,'T_Pyruvate',0,'u');
% turn off lactate uptake
model = changeRxnBounds(model,'T_Lactate',0,'l');
% turn off CO2 uptake
model = changeRxnBounds(model,'T_CO2',0,'l');
% turn off HCO3 uptake
model = changeRxnBounds(model,'T_HCO3',1,'u');
% have to force transhydrogenase in the forward direction for the
% predictions to run properly. Otherwise, there seems to be an issue with
% Cobratoolbox predictions and ATP production is not actually maximized
model = changeRxnBounds(model,'TransHyd',0,'b');
% prevent glucose uptake
model = changeRxnBounds(model,'T_Glucose_1',0,'b');
model = changeRxnBounds(model,'T_Glucose_2',0,'b');
% turn off glutamine transporter that does not require sodium
model = changeRxnBounds(model,'T_Glutamine_1',0,'b');
% prevent synthesis of collagen VI and collagen II
model = changeRxnBounds(model,'CollagenVI',0,'b');
model = changeRxnBounds(model,'CollagenII',0,'b');
% prevent use of aerobic electron transport chain
model = changeRxnBounds(model,'ETC_4',0,'b');
% set maximization criteria to ATPm
model = changeObjective(model,'T_ATP');
% maximize production of collagen
FBAsolution3 = optimizeCbModel(model,'max');
% pull the important values
yield_ATPm_glutamineNa_noETC4 =((FBAsolution3.v(ATPm))./(FBAsolution3.v(glutamine_Na)));

% no O2 as terminal electron acceptor; glucose transporter with no cost
model = readCbModel('replace_with_xml_file_path','fileType','SBML')
% % prevent the export of nonessential amino acids; aren't needed for this
model = changeRxnBounds(model,'cost_alanine',0,'b');
model = changeRxnBounds(model,'cost_arginine',0,'b');
model = changeRxnBounds(model,'cost_asparagine',0,'b');
model = changeRxnBounds(model,'cost_aspartate',0,'b');
model = changeRxnBounds(model,'cost_cysteine',0,'b');
model = changeRxnBounds(model,'cost_glutamate',0,'b');
model = changeRxnBounds(model,'cost_glutamine',0,'b');
model = changeRxnBounds(model,'cost_glycine',0,'b');
model = changeRxnBounds(model,'cost_proline',0,'b');
model = changeRxnBounds(model,'cost_serine',0,'b');
model = changeRxnBounds(model,'cost_tyrosine',0,'b');
% turn offGTPm
model = changeRxnBounds(model,'T_GTP',0,'b');
% turn off pyruvate uptake
model = changeRxnBounds(model,'T_Pyruvate',0,'u');
% turn off lactate uptake
model = changeRxnBounds(model,'T_Lactate',0,'l');
% turn off CO2 uptake
model = changeRxnBounds(model,'T_CO2',0,'l');
% turn off HCO3 uptake
model = changeRxnBounds(model,'T_HCO3',1,'u');
% have to force transhydrogenase in the forward direction for the
% predictions to run properly. Otherwise, there seems to be an issue with
% Cobratoolbox predictions and ATP production is not actually maximized
model = changeRxnBounds(model,'TransHyd',0,'b');
% prevent glucose uptake
model = changeRxnBounds(model,'T_Glucose_1',0,'b');
model = changeRxnBounds(model,'T_Glucose_2',0,'b');
% turn off glutamine transporter that requires sodium
model = changeRxnBounds(model,'T_Glutamine_2',0,'b');
% prevent use of O2 as terminal electron acceptor
model = changeRxnBounds(model,'ETC_4',0,'b');
% prevent synthesis of collagen VI and collagen II
model = changeRxnBounds(model,'CollagenVI',0,'b');
model = changeRxnBounds(model,'CollagenII',0,'b');
% set maximization criteria to ATPm
model = changeObjective(model,'T_ATP');
% maximize production of collagen
FBAsolution4 = optimizeCbModel(model,'max');
% pull the important values
yield_ATPm_glutaminenoCost_noETC4 =((FBAsolution4.v(ATPm))./(FBAsolution4.v(glutamine_noCost)));


%% Model predictions for collagen II yields on glutamine 

% first, start with aerobic conditions, use the glutamine transporter that
% requires Na+
model = readCbModel('replace_with_xml_file_path','fileType','SBML')
% % prevent the export of nonessential amino acids; aren't needed for this
model = changeRxnBounds(model,'cost_alanine',0,'b');
model = changeRxnBounds(model,'cost_arginine',0,'b');
model = changeRxnBounds(model,'cost_asparagine',0,'b');
model = changeRxnBounds(model,'cost_aspartate',0,'b');
model = changeRxnBounds(model,'cost_cysteine',0,'b');
model = changeRxnBounds(model,'cost_glutamate',0,'b');
model = changeRxnBounds(model,'cost_glutamine',0,'b');
model = changeRxnBounds(model,'cost_glycine',0,'b');
model = changeRxnBounds(model,'cost_proline',0,'b');
model = changeRxnBounds(model,'cost_serine',0,'b');
model = changeRxnBounds(model,'cost_tyrosine',0,'b');
% turn off ATPm and GTPm
model = changeRxnBounds(model,'T_ATP',0,'b');
model = changeRxnBounds(model,'T_GTP',0,'b');
% turn off pyruvate uptake
model = changeRxnBounds(model,'T_Pyruvate',0,'u');
% turn off lactate uptake
model = changeRxnBounds(model,'T_Lactate',0,'l');
% turn off CO2 uptake
model = changeRxnBounds(model,'T_CO2',0,'l');
% turn off HCO3 uptake
model = changeRxnBounds(model,'T_HCO3',1,'u');
% have to force transhydrogenase in the forward direction for the
% predictions to run properly. Otherwise, there seems to be an issue with
% Cobratoolbox predictions and ATP production is not actually maximized
model = changeRxnBounds(model,'TransHyd',0,'b');
% prevent glucose uptake
model = changeRxnBounds(model,'T_Glucose_1',0,'b');
model = changeRxnBounds(model,'T_Glucose_2',0,'b');
% turn off glutamine transporter that does not require sodium
model = changeRxnBounds(model,'T_Glutamine_1',0,'b');
% prevent synthesis of collagen VI
model = changeRxnBounds(model,'CollagenVI',0,'b');
% set maximization criteria to collagen II
model = changeObjective(model,'T_Collagen_II');
% maximize production of collagen
FBAsolution1 = optimizeCbModel(model,'max');
% pull the important values
yield_CollagenII_glutamineNa =((FBAsolution1.v(collagenII)*Cmol_collagenII)./(FBAsolution1.v(glutamine_Na)*Cmol_glutamine));

% next, aerobic conditions, use the transporter with no cost
model = readCbModel('replace_with_xml_file_path','fileType','SBML')
% prevent the export of nonessential amino acids; aren't needed for this
model = changeRxnBounds(model,'cost_alanine',0,'b');
model = changeRxnBounds(model,'cost_arginine',0,'b');
model = changeRxnBounds(model,'cost_asparagine',0,'b');
model = changeRxnBounds(model,'cost_aspartate',0,'b');
model = changeRxnBounds(model,'cost_cysteine',0,'b');
model = changeRxnBounds(model,'cost_glutamate',0,'b');
model = changeRxnBounds(model,'cost_glutamine',0,'b');
model = changeRxnBounds(model,'cost_glycine',0,'b');
model = changeRxnBounds(model,'cost_proline',0,'b');
model = changeRxnBounds(model,'cost_serine',0,'b');
model = changeRxnBounds(model,'cost_tyrosine',0,'b');
% turn off ATPm and GTPm
model = changeRxnBounds(model,'T_ATP',0,'b');
model = changeRxnBounds(model,'T_GTP',0,'b');
% turn off pyruvate uptake
model = changeRxnBounds(model,'T_Pyruvate',0,'u');
% turn off lactate uptake
model = changeRxnBounds(model,'T_Lactate',0,'l');
% turn off CO2 uptake
model = changeRxnBounds(model,'T_CO2',0,'l');
% turn off HCO3 uptake
model = changeRxnBounds(model,'T_HCO3',1,'u');
% have to force transhydrogenase in the forward direction for the
% predictions to run properly. Otherwise, there seems to be an issue with
% Cobratoolbox predictions and ATP production is not actually maximized
model = changeRxnBounds(model,'TransHyd',0,'b');
% prevent glucose uptake
model = changeRxnBounds(model,'T_Glucose_1',0,'b');
model = changeRxnBounds(model,'T_Glucose_2',0,'b');
% turn off glutamine transporter that requires sodium
model = changeRxnBounds(model,'T_Glutamine_2',0,'b');
% prevent synthesis of collagen VI
model = changeRxnBounds(model,'CollagenVI',0,'b');
% set maximization criteria to collagen II
model = changeObjective(model,'T_Collagen_II');
% maximize production of collagen
FBAsolution2 = optimizeCbModel(model,'max');
% pull the important values
yield_CollagenII_glutaminenoCost =((FBAsolution2.v(collagenII)*Cmol_collagenII)./(FBAsolution2.v(glutamine_noCost)*Cmol_glutamine));

% no O2 as terminal electron acceptor, use the glucose transporter that requires Na+
model = readCbModel('replace_with_xml_file_path','fileType','SBML')
% prevent the export of nonessential amino acids; aren't needed for this
model = changeRxnBounds(model,'cost_alanine',0,'b');
model = changeRxnBounds(model,'cost_arginine',0,'b');
model = changeRxnBounds(model,'cost_asparagine',0,'b');
model = changeRxnBounds(model,'cost_aspartate',0,'b');
model = changeRxnBounds(model,'cost_cysteine',0,'b');
model = changeRxnBounds(model,'cost_glutamate',0,'b');
model = changeRxnBounds(model,'cost_glutamine',0,'b');
model = changeRxnBounds(model,'cost_glycine',0,'b');
model = changeRxnBounds(model,'cost_proline',0,'b');
model = changeRxnBounds(model,'cost_serine',0,'b');
model = changeRxnBounds(model,'cost_tyrosine',0,'b');
% turn off ATPm and GTPm
model = changeRxnBounds(model,'T_ATP',0,'b');
model = changeRxnBounds(model,'T_GTP',0,'b');
% turn off pyruvate uptake
model = changeRxnBounds(model,'T_Pyruvate',0,'u');
% turn off lactate uptake
model = changeRxnBounds(model,'T_Lactate',0,'l');
% turn off CO2 uptake
model = changeRxnBounds(model,'T_CO2',0,'l');
% turn off HCO3 uptake
model = changeRxnBounds(model,'T_HCO3',1,'u');
% have to force transhydrogenase in the forward direction for the
% predictions to run properly. Otherwise, there seems to be an issue with
% Cobratoolbox predictions and ATP production is not actually maximized
model = changeRxnBounds(model,'TransHyd',0,'b');
% prevent glutamine uptake
model = changeRxnBounds(model,'T_Glucose_1',0,'b');
model = changeRxnBounds(model,'T_Glucose_2',0,'b');
% turn off glucose transporter that does not require sodium
model = changeRxnBounds(model,'T_Glutamine_1',0,'b');
% prevent synthesis of collagen VI
model = changeRxnBounds(model,'CollagenVI',0,'b');
% prevent use of O2 as terminal electron acceptor
model = changeRxnBounds(model,'ETC_4',0,'b');
% set maximization criteria to collagen II
model = changeObjective(model,'T_Collagen_II');
% maximize production of collagen
FBAsolution3 = optimizeCbModel(model,'max');
% pull the important values
yield_CollagenII_glutamineNa_noETC4 =((FBAsolution3.v(collagenII)*Cmol_collagenII)./(FBAsolution3.v(glutamine_Na)*Cmol_glutamine));

% no O2 as terminal electron acceptor, use the no cost glucose transporter
model = readCbModel('replace_with_xml_file_path','fileType','SBML')
% prevent the export of nonessential amino acids; aren't needed for this
model = changeRxnBounds(model,'cost_alanine',0,'b');
model = changeRxnBounds(model,'cost_arginine',0,'b');
model = changeRxnBounds(model,'cost_asparagine',0,'b');
model = changeRxnBounds(model,'cost_aspartate',0,'b');
model = changeRxnBounds(model,'cost_cysteine',0,'b');
model = changeRxnBounds(model,'cost_glutamate',0,'b');
model = changeRxnBounds(model,'cost_glutamine',0,'b');
model = changeRxnBounds(model,'cost_glycine',0,'b');
model = changeRxnBounds(model,'cost_proline',0,'b');
model = changeRxnBounds(model,'cost_serine',0,'b');
model = changeRxnBounds(model,'cost_tyrosine',0,'b');
% turn off ATPm and GTPm
model = changeRxnBounds(model,'T_ATP',0,'b');
model = changeRxnBounds(model,'T_GTP',0,'b');
% turn off pyruvate uptake
model = changeRxnBounds(model,'T_Pyruvate',0,'u');
% turn off lactate uptake
model = changeRxnBounds(model,'T_Lactate',0,'l');
% turn off CO2 uptake
model = changeRxnBounds(model,'T_CO2',0,'l');
% turn off HCO3 uptake
model = changeRxnBounds(model,'T_HCO3',1,'u');
% have to force transhydrogenase in the forward direction for the
% predictions to run properly. Otherwise, there seems to be an issue with
% Cobratoolbox predictions and ATP production is not actually maximized
model = changeRxnBounds(model,'TransHyd',0,'b');
% prevent glucose uptake
model = changeRxnBounds(model,'T_Glucose_1',0,'b');
model = changeRxnBounds(model,'T_Glucose_2',0,'b');
% turn off glutamine transporter that does not require sodium
model = changeRxnBounds(model,'T_Glutamine_2',0,'b');
% prevent synthesis of collagen VI
model = changeRxnBounds(model,'CollagenVI',0,'b');
% prevent use of O2 as terminal electron acceptor
model = changeRxnBounds(model,'ETC_4',0,'b');
% set maximization criteria to collagen II
model = changeObjective(model,'T_Collagen_II');
% maximize production of collagen
FBAsolution4 = optimizeCbModel(model,'max');
% pull the important values
yield_CollagenII_glutaminenoCost_noETC4 =((FBAsolution4.v(collagenII)*Cmol_collagenII)./(FBAsolution4.v(glutamine_noCost)*Cmol_glutamine));

%% Model predictions for collagen VI yields on glutamine 

% first, start with aerobic conditions, use the glutamine transporter that
% requires Na+
model = readCbModel('replace_with_xml_file_path','fileType','SBML')
% % prevent the export of nonessential amino acids; aren't needed for this
model = changeRxnBounds(model,'cost_alanine',0,'b');
model = changeRxnBounds(model,'cost_arginine',0,'b');
model = changeRxnBounds(model,'cost_asparagine',0,'b');
model = changeRxnBounds(model,'cost_aspartate',0,'b');
model = changeRxnBounds(model,'cost_cysteine',0,'b');
model = changeRxnBounds(model,'cost_glutamate',0,'b');
model = changeRxnBounds(model,'cost_glutamine',0,'b');
model = changeRxnBounds(model,'cost_glycine',0,'b');
model = changeRxnBounds(model,'cost_proline',0,'b');
model = changeRxnBounds(model,'cost_serine',0,'b');
model = changeRxnBounds(model,'cost_tyrosine',0,'b');
% turn off ATPm and GTPm
model = changeRxnBounds(model,'T_ATP',0,'b');
model = changeRxnBounds(model,'T_GTP',0,'b');
% turn off pyruvate uptake
model = changeRxnBounds(model,'T_Pyruvate',0,'u');
% turn off lactate uptake
model = changeRxnBounds(model,'T_Lactate',0,'l');
% turn off CO2 uptake
model = changeRxnBounds(model,'T_CO2',0,'l');
% turn off HCO3 uptake
model = changeRxnBounds(model,'T_HCO3',1,'u');
% have to force transhydrogenase in the forward direction for the
% predictions to run properly. Otherwise, there seems to be an issue with
% Cobratoolbox predictions and ATP production is not actually maximized
model = changeRxnBounds(model,'TransHyd',0,'b');
% prevent glucose uptake
model = changeRxnBounds(model,'T_Glucose_1',0,'b');
model = changeRxnBounds(model,'T_Glucose_2',0,'b');
% turn off glutamine transporter that does not require sodium
model = changeRxnBounds(model,'T_Glutamine_1',0,'b');
% prevent synthesis of collagen II
model = changeRxnBounds(model,'CollagenII',0,'b');
% set maximization criteria to collagen VI
model = changeObjective(model,'T_Collagen_VI');
% maximize production of collagen
FBAsolution1 = optimizeCbModel(model,'max');
% pull the important values
yield_CollagenVI_glutamineNa =((FBAsolution1.v(collagenVI)*Cmol_collagenVI)./(FBAsolution1.v(glutamine_Na)*Cmol_glutamine));

% next, aerobic conditions, use the transporter with no cost
model = readCbModel('replace_with_xml_file_path','fileType','SBML')
% prevent the export of nonessential amino acids; aren't needed for this
model = changeRxnBounds(model,'cost_alanine',0,'b');
model = changeRxnBounds(model,'cost_arginine',0,'b');
model = changeRxnBounds(model,'cost_asparagine',0,'b');
model = changeRxnBounds(model,'cost_aspartate',0,'b');
model = changeRxnBounds(model,'cost_cysteine',0,'b');
model = changeRxnBounds(model,'cost_glutamate',0,'b');
model = changeRxnBounds(model,'cost_glutamine',0,'b');
model = changeRxnBounds(model,'cost_glycine',0,'b');
model = changeRxnBounds(model,'cost_proline',0,'b');
model = changeRxnBounds(model,'cost_serine',0,'b');
model = changeRxnBounds(model,'cost_tyrosine',0,'b');
% turn off ATPm and GTPm
model = changeRxnBounds(model,'T_ATP',0,'b');
model = changeRxnBounds(model,'T_GTP',0,'b');
% turn off pyruvate uptake
model = changeRxnBounds(model,'T_Pyruvate',0,'u');
% turn off lactate uptake
model = changeRxnBounds(model,'T_Lactate',0,'l');
% turn off CO2 uptake
model = changeRxnBounds(model,'T_CO2',0,'l');
% turn off HCO3 uptake
model = changeRxnBounds(model,'T_HCO3',1,'u');
% have to force transhydrogenase in the forward direction for the
% predictions to run properly. Otherwise, there seems to be an issue with
% Cobratoolbox predictions and ATP production is not actually maximized
model = changeRxnBounds(model,'TransHyd',0,'b');
% prevent glucose uptake
model = changeRxnBounds(model,'T_Glucose_1',0,'b');
model = changeRxnBounds(model,'T_Glucose_2',0,'b');
% turn off glutamine transporter that requires sodium
model = changeRxnBounds(model,'T_Glutamine_2',0,'b');
% prevent synthesis of collagen II
model = changeRxnBounds(model,'CollagenII',0,'b');
% set maximization criteria to collagen VI
model = changeObjective(model,'T_Collagen_VI');
% maximize production of collagen
FBAsolution2 = optimizeCbModel(model,'max');
% pull the important values
yield_CollagenVI_glutaminenoCost =((FBAsolution2.v(collagenVI)*Cmol_collagenVI)./(FBAsolution2.v(glutamine_noCost)*Cmol_glutamine));

% no O2 as terminal electron acceptor, use the glucose transporter that requires Na+
model = readCbModel('replace_with_xml_file_path','fileType','SBML')
% prevent the export of nonessential amino acids; aren't needed for this
model = changeRxnBounds(model,'cost_alanine',0,'b');
model = changeRxnBounds(model,'cost_arginine',0,'b');
model = changeRxnBounds(model,'cost_asparagine',0,'b');
model = changeRxnBounds(model,'cost_aspartate',0,'b');
model = changeRxnBounds(model,'cost_cysteine',0,'b');
model = changeRxnBounds(model,'cost_glutamate',0,'b');
model = changeRxnBounds(model,'cost_glutamine',0,'b');
model = changeRxnBounds(model,'cost_glycine',0,'b');
model = changeRxnBounds(model,'cost_proline',0,'b');
model = changeRxnBounds(model,'cost_serine',0,'b');
model = changeRxnBounds(model,'cost_tyrosine',0,'b');
% turn off ATPm and GTPm
model = changeRxnBounds(model,'T_ATP',0,'b');
model = changeRxnBounds(model,'T_GTP',0,'b');
% turn off pyruvate uptake
model = changeRxnBounds(model,'T_Pyruvate',0,'u');
% turn off lactate uptake
model = changeRxnBounds(model,'T_Lactate',0,'l');
% turn off CO2 uptake
model = changeRxnBounds(model,'T_CO2',0,'l');
% turn off HCO3 uptake
model = changeRxnBounds(model,'T_HCO3',1,'u');
% have to force transhydrogenase in the forward direction for the
% predictions to run properly. Otherwise, there seems to be an issue with
% Cobratoolbox predictions and ATP production is not actually maximized
model = changeRxnBounds(model,'TransHyd',0,'b');
% prevent glutamine uptake
model = changeRxnBounds(model,'T_Glucose_1',0,'b');
model = changeRxnBounds(model,'T_Glucose_2',0,'b');
% turn off glucose transporter that does not require sodium
model = changeRxnBounds(model,'T_Glutamine_1',0,'b');
% prevent use of O2 as terminal electron acceptor
model = changeRxnBounds(model,'ETC_4',0,'b');
% prevent synthesis of collagen II
model = changeRxnBounds(model,'CollagenII',0,'b');
% set maximization criteria to collagen VI
model = changeObjective(model,'T_Collagen_VI');
% maximize production of collagen
FBAsolution3 = optimizeCbModel(model,'max');
% pull the important values
yield_CollagenVI_glutamineNa_noETC4 =((FBAsolution3.v(collagenVI)*Cmol_collagenVI)./(FBAsolution3.v(glutamine_Na)*Cmol_glutamine));

% no O2 as terminal electron acceptor, use the no cost glucose transporter
model = readCbModel('replace_with_xml_file_path','fileType','SBML')
% prevent the export of nonessential amino acids; aren't needed for this
model = changeRxnBounds(model,'cost_alanine',0,'b');
model = changeRxnBounds(model,'cost_arginine',0,'b');
model = changeRxnBounds(model,'cost_asparagine',0,'b');
model = changeRxnBounds(model,'cost_aspartate',0,'b');
model = changeRxnBounds(model,'cost_cysteine',0,'b');
model = changeRxnBounds(model,'cost_glutamate',0,'b');
model = changeRxnBounds(model,'cost_glutamine',0,'b');
model = changeRxnBounds(model,'cost_glycine',0,'b');
model = changeRxnBounds(model,'cost_proline',0,'b');
model = changeRxnBounds(model,'cost_serine',0,'b');
model = changeRxnBounds(model,'cost_tyrosine',0,'b');
% turn off ATPm and GTPm
model = changeRxnBounds(model,'T_ATP',0,'b');
model = changeRxnBounds(model,'T_GTP',0,'b');
% turn off pyruvate uptake
model = changeRxnBounds(model,'T_Pyruvate',0,'u');
% turn off lactate uptake
model = changeRxnBounds(model,'T_Lactate',0,'l');
% turn off CO2 uptake
model = changeRxnBounds(model,'T_CO2',0,'l');
% turn off HCO3 uptake
model = changeRxnBounds(model,'T_HCO3',1,'u');
% have to force transhydrogenase in the forward direction for the
% predictions to run properly. Otherwise, there seems to be an issue with
% Cobratoolbox predictions and ATP production is not actually maximized
model = changeRxnBounds(model,'TransHyd',0,'b');
% prevent glucose uptake
model = changeRxnBounds(model,'T_Glucose_1',0,'b');
model = changeRxnBounds(model,'T_Glucose_2',0,'b');
% turn off glutamine transporter that does not require sodium
model = changeRxnBounds(model,'T_Glutamine_2',0,'b');
% prevent use of O2 as terminal electron acceptor
model = changeRxnBounds(model,'ETC_4',0,'b');
% prevent synthesis of collagen II
model = changeRxnBounds(model,'CollagenII',0,'b');
% set maximization criteria to collagen VI
model = changeObjective(model,'T_Collagen_VI');
% maximize production of collagen
FBAsolution4 = optimizeCbModel(model,'max');
% pull the important values
yield_CollagenVI_glutaminenoCost_noETC4 =((FBAsolution4.v(collagenVI)*Cmol_collagenVI)./(FBAsolution4.v(glutamine_noCost)*Cmol_glutamine));

%% AMINO ACID YIELDS ON GLUCOSE, AEROBIC
% predict yields of individual amino acids on glucose. Use the glucose
% transporter that requires Na+. Aerobic conditions.
model = readCbModel('replace_with_xml_file_path','fileType','SBML')
 
% turn off ATPm and GTPm
model = changeRxnBounds(model,'T_GTP',0,'b');
model = changeRxnBounds(model,'T_ATP',0,'b');
% turn off pyruvate uptake
model = changeRxnBounds(model,'T_Pyruvate',0,'u');
% turn off lactate uptake
model = changeRxnBounds(model,'T_Lactate',0,'l');
% turn off CO2 uptake
model = changeRxnBounds(model,'T_CO2',0,'l');
% turn off HCO3 uptake
model = changeRxnBounds(model,'T_HCO3',1,'u');
% cysteine production is causing loops that result in CO2 fixation...turn
% it off
model = changeRxnBounds(model,'cost_cysteine',0,'b');
% have to force transhydrogenase in the forward direction for the
% predictions to run properly. Otherwise, there seems to be an issue with
% Cobratoolbox predictions and ATP production is not actually maximized
model = changeRxnBounds(model,'TransHyd',0,'b');
% prevent glutamine uptake
model = changeRxnBounds(model,'T_Glutamine_1',0,'b');
model = changeRxnBounds(model,'T_Glutamine_2',0,'b');
% turn off glucose transporter that does not require sodium
model = changeRxnBounds(model,'T_Glucose_1',0,'b');
% prevent synthesis of collagen VI and collagen II
model = changeRxnBounds(model,'CollagenVI',0,'b');
model = changeRxnBounds(model,'CollagenII',0,'b');
% set maximization criteria to the amino acid
model = changeObjective(model,'cost_alanine');
% maximize production of objective
FBAsolution_alanine = optimizeCbModel(model,'max');
% pull the important values
yield_alanine_glucoseNa =((FBAsolution_alanine.v(alanine)*Cmol_alanine)./(FBAsolution_alanine.v(glucose_Na)*Cmol_glucose));


% set maximization criteria to the amino acid
model = changeObjective(model,'cost_arginine');
% maximize production of objective
FBAsolution_arginine = optimizeCbModel(model,'max');
% pull the important values
yield_arginine_glucoseNa =((FBAsolution_arginine.v(arginine)*Cmol_arginine)./(FBAsolution_arginine.v(glucose_Na)*Cmol_glucose));


% set maximization criteria to the amino acid
model = changeObjective(model,'cost_asparagine');
% maximize production of objective
FBAsolution_asparagine = optimizeCbModel(model,'max');
% pull the important values
yield_asparagine_glucoseNa =((FBAsolution_asparagine.v(asparagine)*Cmol_asparagine)./(FBAsolution_asparagine.v(glucose_Na)*Cmol_glucose));


% set maximization criteria to the amino acid
model = changeObjective(model,'cost_aspartate');
% maximize production of objective
FBAsolution_aspartate = optimizeCbModel(model,'max');
% pull the important values
yield_aspartate_glucoseNa =((FBAsolution_aspartate.v(aspartate)*Cmol_aspartate)./(FBAsolution_aspartate.v(glucose_Na)*Cmol_glucose));

% set maximization criteria to the amino acid
%getting an infinite yield still.. limit bicarb uptake at some point
model = changeRxnBounds(model,'cost_cysteine',100,'u');
model = changeObjective(model,'cost_cysteine');
% maximize production of objective
FBAsolution_cysteine = optimizeCbModel(model,'max');
% pull the important values
yield_cysteine_glucoseNa =((FBAsolution_cysteine.v(cysteine)*Cmol_cysteine)./(FBAsolution_cysteine.v(glucose_Na)*Cmol_glucose));


% cysteine production is causing loops that result in CO2 fixation...turn
% it off
model = changeRxnBounds(model,'cost_cysteine',0,'b');

% set maximization criteria to the amino acid
model = changeObjective(model,'cost_glutamate');
% maximize production of objective
FBAsolution_glutamate = optimizeCbModel(model,'max');
% pull the important values
yield_glutamate_glucoseNa =((FBAsolution_glutamate.v(glutamate)*Cmol_glutamate)./(FBAsolution_glutamate.v(glucose_Na)*Cmol_glucose));

% set maximization criteria to the amino acid
model = changeObjective(model,'cost_glutamine');
% maximize production of objective
FBAsolution_glutamine = optimizeCbModel(model,'max');
% pull the important values
yield_glutamine_glucoseNa =((FBAsolution_glutamine.v(glutamine)*Cmol_glutamine)./(FBAsolution_glutamine.v(glucose_Na)*Cmol_glucose));

% set maximization criteria to the amino acid
model = changeObjective(model,'cost_glycine');
% maximize production of objective
FBAsolution_glycine = optimizeCbModel(model,'max');
% pull the important values
yield_glycine_glucoseNa =((FBAsolution_glycine.v(glycine)*Cmol_glycine)./(FBAsolution_glycine.v(glucose_Na)*Cmol_glucose));

% set maximization criteria to the amino acid
model = changeObjective(model,'cost_proline');
% maximize production of objective
FBAsolution_proline = optimizeCbModel(model,'max');
% pull the important values
yield_proline_glucoseNa =((FBAsolution_proline.v(proline)*Cmol_proline)./(FBAsolution_proline.v(glucose_Na)*Cmol_glucose));

% set maximization criteria to the amino acid
model = changeObjective(model,'cost_serine');
% maximize production of objective
FBAsolution_serine = optimizeCbModel(model,'max');
% pull the important values
yield_serine_glucoseNa =((FBAsolution_serine.v(serine)*Cmol_serine)./(FBAsolution_serine.v(glucose_Na)*Cmol_glucose));

% set maximization criteria to the amino acid
model = changeObjective(model,'cost_tyrosine');
% maximize production of objective
FBAsolution_tyrosine = optimizeCbModel(model,'max');
% pull the important values
yield_tyrosine_glucoseNa =((FBAsolution_tyrosine.v(tyrosine)*Cmol_tyrosine)./(FBAsolution_tyrosine.v(glucose_Na)*Cmol_glucose));
% pull the important values
yield_tyrosine_phenylalanine =((FBAsolution_tyrosine.v(tyrosine)*Cmol_tyrosine)./(FBAsolution_tyrosine.v(phenylalanine)*Cmol_phenylalanine));


%% AMINO ACID YIELDS ON GLUCOSE, NO OXIDATIVE PHOSPHORYLATION
% predict yields of individual amino acids on glucose. Use the glucose
% transporter that requires Na+. Turn off ETC4 (terminal oxidase). These
% predictions are not anaerobic exactly-tyrosine production requires O2
model = readCbModel('replace_with_xml_file_path','fileType','SBML')
% 
% turn off ATPm and GTPm
model = changeRxnBounds(model,'T_GTP',0,'b');
model = changeRxnBounds(model,'T_ATP',0,'b');
% turn off pyruvate uptake
model = changeRxnBounds(model,'T_Pyruvate',0,'u');
% turn off lactate uptake
model = changeRxnBounds(model,'T_Lactate',0,'l');
% turn off CO2 uptake
model = changeRxnBounds(model,'T_CO2',0,'l');
% turn off HCO3 uptake
model = changeRxnBounds(model,'T_HCO3',1,'u');
% have to force transhydrogenase in the forward direction for the
% predictions to run properly. Otherwise, there seems to be an issue with
% Cobratoolbox predictions and ATP production is not actually maximized
model = changeRxnBounds(model,'TransHyd',0,'b');
% prevent glutamine uptake
model = changeRxnBounds(model,'T_Glutamine_1',0,'b');
model = changeRxnBounds(model,'T_Glutamine_2',0,'b');
% turn off glucose transporter that does not require sodium
model = changeRxnBounds(model,'T_Glucose_1',0,'b');
% prevent synthesis of collagen VI and collagen II
model = changeRxnBounds(model,'CollagenVI',0,'b');
model = changeRxnBounds(model,'CollagenII',0,'b');
% prevent use of ETC_4
model = changeRxnBounds(model,'ETC_4',0,'b');


% set maximization criteria to the amino acid
model = changeObjective(model,'cost_alanine');
% maximize production of objective
FBAsolution_alanine_noETC4 = optimizeCbModel(model,'max');
% pull the important values
yield_alanine_glucoseNa_noETC4 =((FBAsolution_alanine_noETC4.v(alanine)*Cmol_alanine)./(FBAsolution_alanine_noETC4.v(glucose_Na)*Cmol_glucose));


% set maximization criteria to the amino acid
model = changeObjective(model,'cost_arginine');
% maximize production of objective
FBAsolution_arginine_noETC4 = optimizeCbModel(model,'max');
% pull the important values
yield_arginine_glucoseNa_noETC4 =((FBAsolution_arginine_noETC4.v(arginine)*Cmol_arginine)./(FBAsolution_arginine_noETC4.v(glucose_Na)*Cmol_glucose));


% set maximization criteria to the amino acid
model = changeObjective(model,'cost_asparagine');
% maximize production of objective
FBAsolution_asparagine_noETC4 = optimizeCbModel(model,'max');
% pull the important values
yield_asparagine_glucoseNa_noETC4 =((FBAsolution_asparagine_noETC4.v(asparagine)*Cmol_asparagine)./(FBAsolution_asparagine_noETC4.v(glucose_Na)*Cmol_glucose));

%turn off cost_tyrosine
model = changeRxnBounds(model,'cost_tyrosine',0,'l');
% set maximization criteria to the amino acid
model = changeObjective(model,'cost_aspartate');
% maximize production of objective
FBAsolution_aspartate_noETC4 = optimizeCbModel(model,'max');
% pull the important values
yield_aspartate_glucoseNa_noETC4 =((FBAsolution_aspartate_noETC4.v(aspartate)*Cmol_aspartate)./(FBAsolution_aspartate_noETC4.v(glucose_Na)*Cmol_glucose));

%turn on cost_tyrosine
model = changeRxnBounds(model,'cost_tyrosine',100,'u');
% set maximization criteria to the amino acid
model = changeObjective(model,'cost_cysteine');
% maximize production of objective
FBAsolution_cysteine_noETC4 = optimizeCbModel(model,'max');
% pull the important values
yield_cysteine_glucoseNa_noETC4 =((FBAsolution_cysteine_noETC4.v(cysteine)*Cmol_cysteine)./(FBAsolution_cysteine_noETC4.v(glucose_Na)*Cmol_glucose));

% set maximization criteria to the amino acid
model = changeObjective(model,'cost_glutamate');
% maximize production of objective
FBAsolution_glutamate_noETC4 = optimizeCbModel(model,'max');
% pull the important values
yield_glutamate_glucoseNa_noETC4 =((FBAsolution_glutamate_noETC4.v(glutamate)*Cmol_glutamate)./(FBAsolution_glutamate_noETC4.v(glucose_Na)*Cmol_glucose));

% set maximization criteria to the amino acid
model = changeObjective(model,'cost_glutamine');
% maximize production of objective
FBAsolution_glutamine_noETC4 = optimizeCbModel(model,'max');
% pull the important values
yield_glutamine_glucoseNa_noETC4 =((FBAsolution_glutamine_noETC4.v(glutamine)*Cmol_glutamine)./(FBAsolution_glutamine_noETC4.v(glucose_Na)*Cmol_glucose));

% set maximization criteria to the amino acid
model = changeObjective(model,'cost_glycine');
% maximize production of objective
FBAsolution_glycine_noETC4 = optimizeCbModel(model,'max');
% pull the important values
yield_glycine_glucoseNa_noETC4 =((FBAsolution_glycine_noETC4.v(glycine)*Cmol_glycine)./(FBAsolution_glycine_noETC4.v(glucose_Na)*Cmol_glucose));

% set maximization criteria to the amino acid
model = changeObjective(model,'cost_proline');
% maximize production of objective
FBAsolution_proline_noETC4 = optimizeCbModel(model,'max');
% pull the important values
yield_proline_glucoseNa_noETC4 =((FBAsolution_proline_noETC4.v(proline)*Cmol_proline)./(FBAsolution_proline_noETC4.v(glucose_Na)*Cmol_glucose));

% set maximization criteria to the amino acid
model = changeObjective(model,'cost_serine');
% maximize production of objective
FBAsolution_serine_noETC4 = optimizeCbModel(model,'max');
% pull the important values
yield_serine_glucoseNa_noETC4 =((FBAsolution_serine_noETC4.v(serine)*Cmol_serine)./(FBAsolution_serine_noETC4.v(glucose_Na)*Cmol_glucose));

% set maximization criteria to the amino acid
model = changeObjective(model,'cost_tyrosine');
% maximize production of objective
FBAsolution_tyrosine_noETC4 = optimizeCbModel(model,'max');
% pull the important values
yield_tyrosine_glucoseNa_noETC4 =((FBAsolution_tyrosine_noETC4.v(tyrosine)*Cmol_tyrosine)./(FBAsolution_tyrosine_noETC4.v(glucose_Na)*Cmol_glucose));
% pull the important values
yield_tyrosine_phenylalanine_noETC4 =((FBAsolution_tyrosine_noETC4.v(tyrosine)*Cmol_tyrosine)./(FBAsolution_tyrosine_noETC4.v(phenylalanine)*Cmol_phenylalanine));
```
