## Supplementary material for "Expanded stoichiometric model of chondrocyte metabolism: response to cyclical shear and compressive loading": SI_3.m

```
%% heatmap comparing collagen II yields vs lactate fermentation
% first, run predictions that maximimize lactate production. Pull max
% lactate flux. Use that value as the maximum, scale down lactate
% production, and run predictions for max collagen II as a function of
% lactate fermentation
% for each one, pull the fluxes through key reactions. Then, plot those
% fluxes as a heatmap

initCobraToolbox()

%% Set some reaction names and balance variables
model = readCbModel('/Users/ebruerdogan/Desktop/ADA_Code_July/chondrocyte_model_27july2023_v2.xml','fileType','SBML')

% biomass and ATP maintenance terms
collagenII = find(strcmp('T_Collagen_II',model.rxns));
collagenVI = find(strcmp('T_Collagen_VI',model.rxns));
O2 = find(strcmp('T_O2',model.rxns));
ATPm = find(strcmp('T_ATP',model.rxns));
glucose_noCost = find(strcmp('T_Glucose_1',model.rxns));
glucose_Na = find(strcmp('T_Glucose_2',model.rxns));
glutamine_noCost = find(strcmp('T_Glutamine_1',model.rxns));
glutamine_Na = find(strcmp('T_Glutamine_2',model.rxns));
pyruvate_dehydrogenase = find(strcmp('PYR_1', model.rxns));
oxidase = find(strcmp('ETC_4', model.rxns));
lactate_production = find(strcmp('T_Lactate', model.rxns));
ATP_synthase = find(strcmp('ATP', model.rxns));
pryuvate_transport = find(strcmp('T_Pyruvate',model.rxns));

alanine = find(strcmp('cost_alanine',model.rxns));
arginine = find(strcmp('cost_arginine',model.rxns));
asparagine = find(strcmp('cost_asparagine',model.rxns));
aspartate = find(strcmp('cost_aspartate',model.rxns));
cysteine = find(strcmp('cost_cysteine',model.rxns));
glutamate = find(strcmp('cost_glutamate',model.rxns));
glutamine = find(strcmp('cost_glutamine',model.rxns));
glycine = find(strcmp('cost_glycine',model.rxns));
proline = find(strcmp('cost_proline',model.rxns));
serine = find(strcmp('cost_serine',model.rxns));
tyrosine = find(strcmp('cost_tyrosine',model.rxns));
phenylalanine = find(strcmp('T_Phe',model.rxns));

% carbon content of each metabolite of interest
Cmol_collagenII = 395;
Cmol_collagenVI = 462;
Cmol_glucose = 6;
Cmol_glutamine = 5;

Cmol_alanine = 3;
Cmol_arginine = 6;
Cmol_asparagine = 4;
Cmol_aspartate = 4;
Cmol_cysteine = 3;
Cmol_glutamate = 5;
Cmol_glycine = 2;
Cmol_proline = 5;
Cmol_serine = 3;
Cmol_tyrosine = 9;
Cmol_phenylalanine = 9;


%% run the first prediction that maximizes collagen production on glucose
% use the no cost glucose transporter
% aerobic conditions but use glucose transporter with no cost
model = readCbModel('/Users/ebruerdogan/Desktop/ADA_Code_July/chondrocyte_model_27july2023_v2.xml','fileType','SBML')
% % prevent the export of nonessential amino acids; aren't needed for this
model = changeRxnBounds(model,'cost_alanine',0,'b');
model = changeRxnBounds(model,'cost_arginine',0,'b');
model = changeRxnBounds(model,'cost_asparagine',0,'b');
model = changeRxnBounds(model,'cost_aspartate',0,'b');
model = changeRxnBounds(model,'cost_cysteine',0,'b');
model = changeRxnBounds(model,'cost_glutamate',0,'b');
model = changeRxnBounds(model,'cost_glutamine',0,'b');
model = changeRxnBounds(model,'cost_glycine',0,'b');
model = changeRxnBounds(model,'cost_proline',0,'b');
model = changeRxnBounds(model,'cost_serine',0,'b');
model = changeRxnBounds(model,'cost_tyrosine',0,'b');
% turn offGTPm
model = changeRxnBounds(model,'T_GTP',0,'b');
% turn off ATPm
model = changeRxnBounds(model,'T_ATP',0,'b');
% turn off pyruvate uptake
model = changeRxnBounds(model,'T_Pyruvate',0,'u');
% turn off lactate uptake
model = changeRxnBounds(model,'T_Lactate',0,'l');
% have to force transhydrogenase in the forward direction for the
% predictions to run properly. Otherwise, there seems to be an issue with
% Cobratoolbox predictions and ATP production is not actually maximized
model = changeRxnBounds(model,'TransHyd',0,'b');
% prevent glutamine uptake
model = changeRxnBounds(model,'T_Glutamine_1',0,'b');
model = changeRxnBounds(model,'T_Glutamine_2',0,'b');
% turn off glucose transporter that requires sodium
model = changeRxnBounds(model,'T_Glucose_2',0,'b');
% prevent synthesis of collagen VI and collagen II
model = changeRxnBounds(model,'CollagenVI',0,'b');
model = changeRxnBounds(model,'CollagenII',0,'b');
% set maximization criteria to lactate
model = changeObjective(model,'T_Lactate');
% maximize production of collagen II
FBAsolution_max1 = optimizeCbModel(model,'max');
% pull the flux of O2 out
max_lactate = FBAsolution_max1.v(lactate_production);

%% now, scale lactate flux to each of ten different values. This one has transhydrogenase forward only
% aerobic conditions but use glucose transporter with no cost
model = readCbModel('/Users/ebruerdogan/Desktop/ADA_Code_July/chondrocyte_model_27july2023_v2.xml','fileType','SBML')
% % prevent the export of nonessential amino acids; aren't needed for this
model = changeRxnBounds(model,'cost_alanine',0,'b');
model = changeRxnBounds(model,'cost_arginine',0,'b');
model = changeRxnBounds(model,'cost_asparagine',0,'b');
model = changeRxnBounds(model,'cost_aspartate',0,'b');
model = changeRxnBounds(model,'cost_cysteine',0,'b');
model = changeRxnBounds(model,'cost_glutamate',0,'b');
model = changeRxnBounds(model,'cost_glutamine',0,'b');
model = changeRxnBounds(model,'cost_glycine',0,'b');
model = changeRxnBounds(model,'cost_proline',0,'b');
model = changeRxnBounds(model,'cost_serine',0,'b');
model = changeRxnBounds(model,'cost_tyrosine',0,'b');
% turn offGTPm
model = changeRxnBounds(model,'T_GTP',0,'b');
% turn off ATPm
model = changeRxnBounds(model,'T_ATP',0,'b');
% turn off pyruvate uptake
model = changeRxnBounds(model,'T_Pyruvate',0,'u');
% turn off lactate uptake
model = changeRxnBounds(model,'T_Lactate',0,'l');
% have to force transhydrogenase in the forward direction for the
% predictions to run properly. Otherwise, there seems to be an issue with
% Cobratoolbox predictions and ATP production is not actually maximized
model = changeRxnBounds(model,'TransHyd',0,'b');
% prevent glutamine uptake
model = changeRxnBounds(model,'T_Glutamine_1',0,'b');
model = changeRxnBounds(model,'T_Glutamine_2',0,'b');
% turn off glucose transporter that requires sodium
model = changeRxnBounds(model,'T_Glucose_2',0,'b');
% prevent synthesis of collagen II
model = changeRxnBounds(model,'CollagenII',0,'b');
% set maximization criteria to Collagen VI
model = changeObjective(model,'CollagenVI');

lactate_values = [max_lactate, 0.9*max_lactate, 0.8*max_lactate, 0.7*max_lactate, 0.6*max_lactate,0.5*max_lactate,0.4*max_lactate,0.3*max_lactate,0.2*max_lactate,0.1*max_lactate,0];
heatmap_matrix = zeros([11,4])
for i = 1:11
    % set lactate flux
    model = changeRxnBounds(model,'T_Lactate',lactate_values(i),'b');
    lactate_values(i)
    % maximize production of collagen II
    FBAsolution = optimizeCbModel(model,'max');
    % get the values for the fluxes through reactions of interest. Scale to
    % glucose uptake
    flux_oxidase =((FBAsolution.v(oxidase))./(FBAsolution.v(glucose_noCost)));
    heatmap_matrix(i,1) = flux_oxidase;
    flux_collagen =((FBAsolution.v(collagenVI))./(FBAsolution.v(glucose_noCost)));
    heatmap_matrix(i,2) = flux_collagen;
    flux_pyruvatedehydrogenase =((FBAsolution.v(pyruvate_dehydrogenase))./(FBAsolution.v(glucose_noCost)));
    heatmap_matrix(i,3) = flux_pyruvatedehydrogenase;
    flux_lactate =((FBAsolution.v(lactate_production))./(FBAsolution.v(glucose_noCost)));
    heatmap_matrix(i,4) = flux_lactate;
end

% plot the data as a heatmap
x_names = ["oxidative phosphorylation"; "collagen production"; 'pyruvate dehydrogenase'; 'lactate'];
%y_label = str(O2_values);
figure(1)
set(gca, 'fontsize', 55)
h = heatmap(x_names, lactate_values, heatmap_matrix)
h.XLabel = "Reactions (mol/mol glucose)"
h.YLabel = "Lactate Fermentation Fluxes (mol lactate / mol glucose)"
h.Title = "Flux as a Function of Lactate Fermentation"
h.ColorScaling = 'scaledcolumns'
```
